# Integrated genetic code expansion and structural bioinformatics reveal disrupted supramolecular assembly in a genetic disorder

**DOI:** 10.1101/2023.07.24.550340

**Authors:** Valerio Marino, Wanchana Phromkrasae, Michele Bertacchi, Paul Cassini, Krittalak Chakrabandhu, Daniele Dell’Orco, Michèle Studer

**Affiliations:** Department of Neurosciences, Biomedicine and Movement Sciences, Section of Biological Chemistry, University of Verona, Verona, Italy; Université Côte d’Azur, CNRS, Inserm, iBV, France

**Keywords:** NR2F1, BBSOAS, pathogenic variants, ligand binding domain (LBD), structural biochemistry, cellular biology, genetic code expansion (GCE), protein stability and affinity, protein interactions, CRABP2

## Abstract

Deciphering the structural effects of variants is essential for understanding the pathophysiological mechanisms of genetic diseases. Using a neurodevelopmental disorder called Bosch-Boonstra-Schaaf Optic Atrophy Syndrome (BBSOAS) as a genetic disease model, we applied a combined Genetic Code Expansion (GCE) and structural bioinformatics strategy to assess the pathogenic impact of several human NR2F1 variants. Nonsense mutations in the ligand binding domain (LBD) resulted in truncated proteins, while missense variants significantly affected the folding of NR2F1 monomers as well as its supramolecular complexes. The GCE-enabled covalent and site-specific capture of transient supramolecular interactions in living cells revealed the variable quaternary conformations of NR2F1 variants and pinpointed the disrupted interplay with dimeric partners and the newly identified cofactor, CRABP2, while the computational analyses of the NR2F1 structure delineated the molecular basis of the impact of the variants on the isolated and complexed structures. The revealed consequence of the pathogenic mutations on the conformation, supramolecular interplay, and alterations in the cell cycle, viability, and subcellular localization of the different variants reflect the heterogeneous disease spectrum and establish the foundation for further understanding the complexity of BBSOAS.

## INTRODUCTION

Untangling the correlation between amino acid mutations in a protein sequence and disease is crucial to understanding protein functional variation and designing effective therapeutic interventions. A missense mutation or truncation in the coding region of a particular gene can lead to structural alterations, which may render the protein non-functional (“loss of function”), or lead to “gain of function” effects, such as functional dysregulation or the formation of toxic aggregates (Dobson, 2003). Mutations located in or near key functional sites are the most likely to affect protein functions. Along with accurate information about protein structure, computational determination of functional residues and cellular experiments can explain the effect and heterogeneity of mutations on proteins. In addition to the prediction of functional sites, several computational tools can be used to calculate free energy changes associated with pathogenetic mutations, thus allowing predictions of whether the mutation will destabilize the structure of the protein, possibly affecting its functions (Capriotti *et al*, 2005; Capriotti *et al*, 2004; Dell’Orco, 2009).

In this study, we focus on NR2F1 (Nuclear Receptor Subfamily 2 Group F Member 1), an evolutionary well-conserved orphan nuclear receptor acting as a strong transcriptional regulator of several genes and playing key roles during embryogenesis with a particular emphasis on the development of the central nervous system (Alfano *et al*, 2013; Tocco *et al*, 2021). Haploinsufficiency of *NR2F1*, due mainly to *de novo* missense/nonsense mutations or whole-gene deletion of only one of the two alleles, leads to an emerging monogenic neurodevelopmental disease, called Bosch–Boonstra–Schaaf Optic Atrophy Syndrome (BBSOAS; OMIM 615722; ORPHA 401777), an autosomal dominant genetic disorder first described in 2014 (Bosch *et al*, 2014). BBSOAS has been so far diagnosed in more than 150 patients worldwide, however, new patients are reported every year, suggesting that the predicted prevalence between 1 in 100,000 to 250,000 people could be an underestimation. BBSOAS symptoms are very heterogeneous in terms of both presence and severity and include optic nerve atrophy (OA) or optic nerve hypoplasia (ONH), cortical/cerebral visual impairment (CVI), moderate to severe intellectual disability (ID), developmental delay (DD), hypotonia, seizures, speech difficulties, motor dysfunctions and autism spectrum disorder (ASD), among others. It is the peculiar combination of these diverse symptoms, particularly CVI and OA, that differentiate BBSOAS patients from those affected by other neurodevelopmental diseases with related features (Bertacchi *et al*, 2022; Chen *et al*, 2016; Rech *et al*, 2020).

Similar symptoms are shared by multiple BBSOAS patients, however, their degree of severity is variable, possibly depending on the location and type of the NR2F1 mutation (Bertacchi *et al*., 2022; Billiet *et al*, 2022; Chen *et al*., 2016; Rech *et al*., 2020). Pathogenic BBSOAS mutations are principally located in the two most conserved functional domains of the protein: the DNA-binding domain (DBD) and the ligand-binding domain (LBD). While the DBD consists of two zinc-finger domains and is responsible for the interaction with either directed, inverted, and everted direct repeats of the consensus sequence (AGGTCA) present in the promoter of target genes (Tang *et al*, 2015), the LBD is predicted to be necessary for protein dimerization and binding to co-regulators, as suggested by the protein structure of other nuclear receptors of the same family (Rastinejad *et al*, 2015). Clinical investigations suggested a genotype-to-phenotype correlation, as BBSOAS patients with reduced protein dosage and functional haploinsufficiency, due to the loss of one copy of the *NR2F1* gene, show a less severe clinical picture than patients with missense point mutations located in the DBD (Bertacchi *et al*., 2022; Bosch *et al*., 2014; Chen *et al*., 2016; Rech *et al*., 2020). Since NR2F1 seems to bind the DNA in the form of homodimers or heterodimers to diverse targets (Cooney *et al*, 1992; Klinge *et al*, 1997; Leng *et al*, 1996; Park *et al*, 2003), point-mutation variants in the DBD will result in dominant-negative effects, in which the mutated form competes for dimerization with the wild-type (WT) protein or with other nuclear receptors of the same family (Tocco *et al*., 2021).

NR2F proteins are named ‘‘orphan’’ receptors since the identity of a ligand binding to the LBD domain is still elusive. Nevertheless, a functionally relevant region of the NR2F1 protein structure is the C-terminal Activation Function 2 helix (named AF2), whose active conformational state, generally obtained *via* interactions with specific ligands, allows the binding of cofactors to the LBD and ultimately controls the transcriptional regulation of target genes (Germain *et al*, 2006). In the specific case of the NR2F1 homolog, NR2F2, crystallographic studies have shown that the LBD is normally present in an auto-inhibited conformation, due to the binding between the AF2 helix and cofactor binding sites, and that this auto-repressed state can be reverted with a high concentration of retinoic acid (RA) (Kruse *et al*, 2008). Due to the high homology between NR2F1 and NR2F2, it is conceivable that a similar mechanism might also be present for NR2F1, but this has not been tested yet. Despite the fact that a genotype-phenotype correlation is starting to emerge (Bertacchi et al., 2022; Rech et al., 2020), it is still unclear whether mutations in the LBD are associated with different clinical symptoms and severity. Furthermore, little is known about the structural function of the LBD and how missense mutations or truncations within this domain specifically impact NR2F1 protein structure and, consequently, cell behavior in pathological conditions.

In this study, we used a multidisciplinary approach ranging from structural biochemistry to cellular biology and genetic code expansion (GCE) to specifically assess the impact of disease-associated LBD variants on the function of the NR2F1 protein. We found that patient-specific LBD pathogenic variants show altered structural stability and abnormal nuclear *versus* cytoplasmic localization, and ultimately impact cell proliferation and survival. Moreover, we found that some NR2F1 variants display altered oligomerization both *in silico* and *in cellula* and, unexpectedly, increased propensity to form homo-/heterodimers or large protein complexes. By using a GCE-enabled covalent and site-specific capturing technique, we assessed the impact of NR2F1 variants on the dimerization and unveiled for the first time an interaction between NR2F1 and CRABP2, which are co-expressed *in vivo* in developing brain structures, such as the eye and the brain. Together, our data shed new light on the impact of BBSOAS LBD mutations on NR2F1 activity and, more in general, on the use of structural analyses and GCE-based approaches to unravel the molecular function of nuclear receptors in both physiological and pathological conditions.

## RESULTS

### Molecular modeling of the NR2F1 LBD in different functional states by homology

NR2F1 is an orphan nuclear receptor belonging to the NR2F subfamily of the nuclear hormone receptor superfamily and thus shares the same structural organization as all other members of the family (**Figure 1**). An N-terminal disordered region is connected to the DNA-binding domain (DBD), which is comprised of two zinc finger motifs arranged in a C4-type domain (residues 84-153) and is connected by a flexible hinge region to the all α-helical ligand-binding domain (LBD, **Figure 1A** and **Video S1**). Currently, no experimental structure of the full-length human NR2F1 is available on the Protein Data Bank (PDB), and the structure predicted by *AlphaFold* is not reliable in either the N-terminal disordered region or the hinge region. By taking advantage of the high sequence identity between NR2F1 and its homolog NR2F2 (97%, **Figure S1**), we modeled the three-dimensional structure of the auto-repressed conformation of the NR2F1 LBD by homology using the experimentally resolved structure of the NR2F2 LBD (PDB entry 3cjw (Kruse et al., 2008),

**Figure 1.**
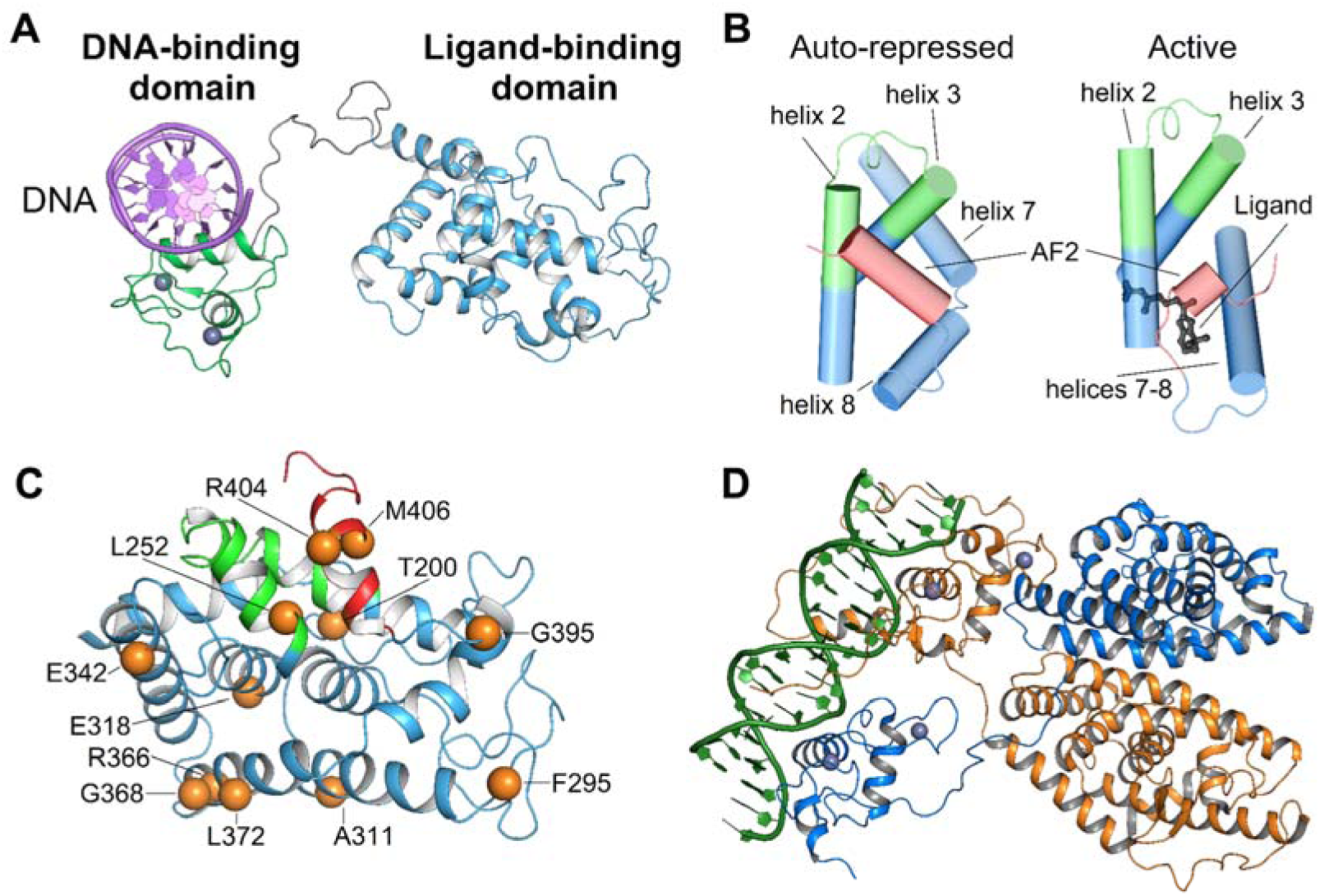
Structural representation of DNA-NR2F1 assembly. (**A**) A three-dimensional structure of NR2F1-DNA complex was modeled by the superimposition of the modeled LBD and DBD-DNA complex on the template structure provided by AlphaFold after removal of residues 1-83 due to unreliable predictions. Protein and DNA structure are represented as a cartoon, with the LBD shown in green, the DBD in cyan, and the dsDNA in purple. (**B**) Conformational changes of the AF2 helix belonging to the LBD upon ligand binding. Protein structures proximal to AF2, namely helices 2, 3, 7, and 8, are shown as blue cylindrical cartoons with AF2 helix displayed in red; the ligand is represented as black sticks with O atoms colored in red, CRS is colored in green. (**C**) The three-dimensional structure of the LBD is shown as a cyan cartoon with the AF2 helix shown in red and the CRS in green. The Cα of the residues whose mutations are associated with BBSOAS are represented by orange spheres. (**D**) Representation of the putative interaction of NR2F1 dimer with DNA based on the homology with RXRα-LXRβ (PDB entry 4NQA) heterodimer. Protein structure is represented as a cartoon with RXRα shown in blue and LXRβ in orange, DNA strands are depicted in green.

The LBD is responsible for the dimerization of nuclear receptors *via* the dimerization interface (DI, (Perlmann *et al*, 1996), **Figure S1**) as well as for the recognition and binding of ligands, co-activators, and co-repressors *via* the co-activator recognition site (CRS, residues 228-253, **Figure 1B**) (Gampe *et al*, 2000; Lu *et al*, 2020). Accessibility of functionally important molecules to LBD is modulated by the conformational switch occurring at the C-terminal region of the protein, which involves the Activation Function 2 helix (AF2, residues 399-408, **Figure 1B**). Indeed, when the LBD is in its auto-repressed conformation, helices 7 and 8 are two separate entities connected by a loop so that the AF2 helix masks the CRS, thus inhibiting potential interactions with co-regulators. Ligand binding to the LBD triggers a structural rearrangement to an active conformation, where the N-terminal part of helix 7 and the C-terminal portion of helix 8 fuse into a single helix that displaces the AF2 helix from the binding pocket, which then becomes available for co-regulators (**Figure 1B**). As in the case with the auto-repressed conformation, no experimental structure of the active form of NR2F1 LBD was available on the PDB. Therefore, we modeled its three-dimensional architecture using as a template the structure of another member of the nuclear receptor superfamily, namely the Retinoid X Receptor RXRα (∼40 % sequence identity with NR2F1 LBD) in complex with 9-cis-retinoic acid (9cRA, PDB entry 1FM6), after removal of the co-activator peptides, as previously described in (Gampe *et al*., 2000; Khalil *et al*, 2022).

### Impact of BBSOAS-associated LBD variants on protein structure

Since missense variants in the LBD have been associated with a variety of BBSOAS symptoms (**Tables 1 and S1**) (Bertacchi *et al*., 2022; Rech *et al*., 2020), we hypothesized that these mutations affect protein structure and function by interfering with the dimerization and binding of the ligand and co-activators. We, therefore, predicted the effects of missense mutations on the folding of both the auto-repressed and the active form of the LBD (**Figure 1C; Tables S2** and **S3**; and **Video S2**) to shed light on the possible genotype-phenotype correlation based on the LBD mutations.

**Table 1.**
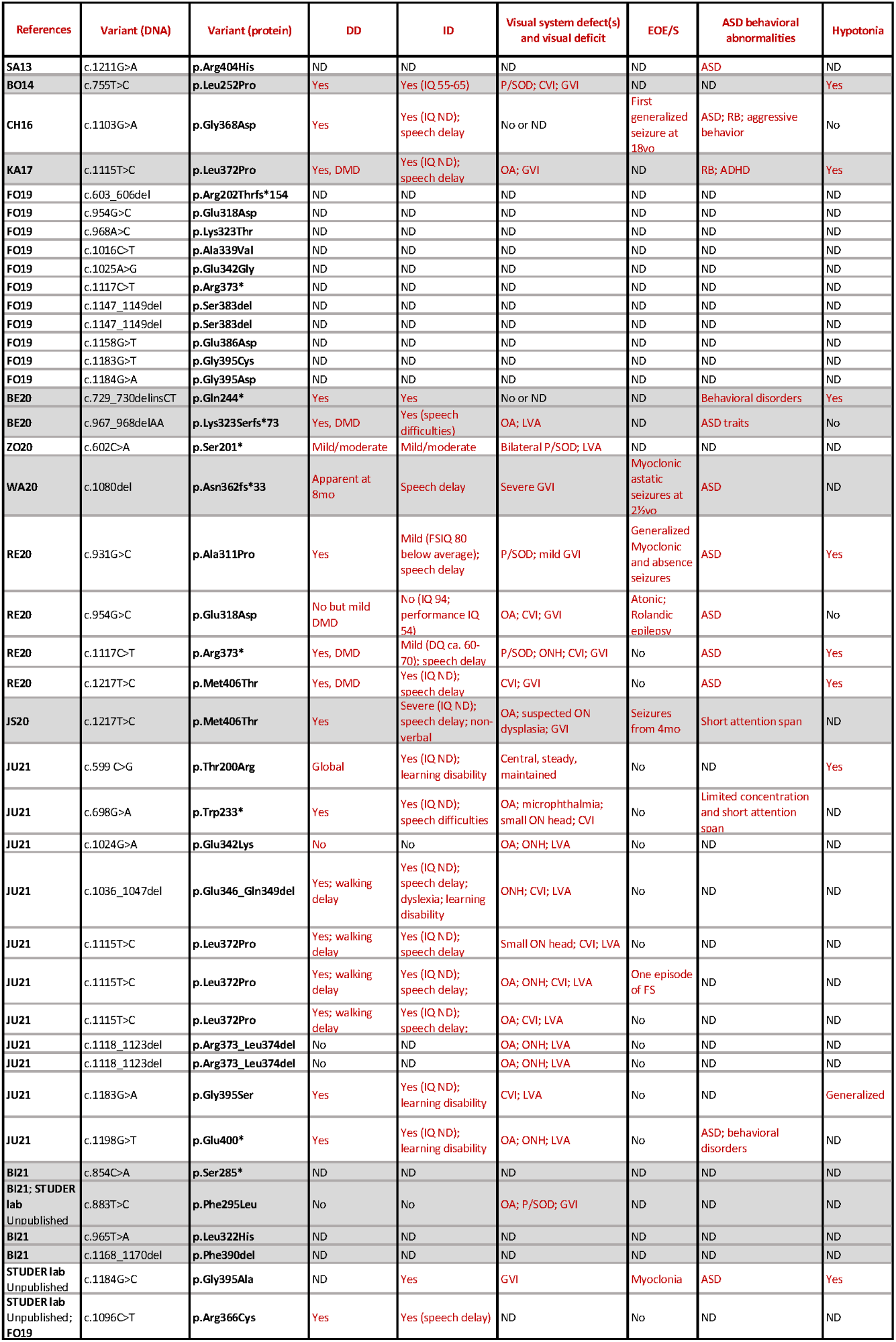
List of known LBD NR2F1 variants and clinical description of corresponding BBSOAS patients. BBSOAS patients carrying LBD variants in their NR2F1 locus, listed following the chronological order of publications describing their cases, and identified by their DNA and protein variant. Main clinical signs (columns) can include developmental delay (DD), intellectual disability (ID), visual system deficits such as optic atrophy (OA) and cortical/cerebral visual impairment (CVI), early-onset epilepsy and seizures (EOE/S), autism spectrum disorder (ASD) and behavioral abnormalities, and hypotonia. An extended version of these data, with additional columns showing the LOVD identifier of NR2F1 variants, clinical data about altered brain morphology as observed by MRI, and other less common clinical features, is available in Table S1. List of references (first column): BE20, (Bertacchi et al, 2020); BO14, (Bosch et al., 2014); CH16, (Chen et al., 2016); FO19, (Fokkema et al, 2019); JS20, (Jezela-Stanek et al, 2020); JU21, (Jurkute et al, 2021); KA17, (Kaiwar et al, 2017); SA13, (Sanders et al, 2012); RE20, (Rech et al., 2020); WA20, (Walsh et al, 2020); ZO20, (Zou et al, 2020). Abbreviations: ADHD, attention deficit hyperactivity disorder; ASD, autism spectrum disorder; DD, developmental delay; DMD, delayed motor development/poor coordination; DQ, developmental quotient; EOE/S, early-onset epilepsy/seizures; FS, febrile seizures; GVI, general visual impairments; ID, intellectual disability; IQ, intelligence quotient; IS, infantile spasms; LBD, ligand binding domain; LVA, low visual acuity; ND, not determined; OA, optic atrophy; ON, optic nerve; ONH, ON hypoplasia; P/SOD, pale/small optic disc; RB, repetitive behavior.

By comparing the calculated relative change in folding free energy (ΔΔG_f_^app^ values, see STAR Methods section, **Table S2**), we found that all but T200R and G368D variants destabilize the folding of the isolated auto-repressed domain, with the largest effect exerted by proline substitutions (L252P, A311P and L372P). Similarly, the same analysis performed on the active form identified the same proline substitution as the most destabilizing, together with the R366C variant (**Table S3**). This was not surprising, since L252 is located on helix 3, whereas A311 and L372 are located on helices 4 and 8, respectively, and proline substitutions are known to disrupt the H-bond pattern required for the proper folding of the α-helix. Moreover, the F295L variant, localized on the long loop connecting helices 3 and 4, was predicted to exert a stabilizing effect on the active form (ΔΔG_f_^app^ = -4.60 kcal/mol) due to a more favorable hydrophobic packing within the protein core. However, upon switching to the auto-repressed form, the same mutation destabilized protein folding (ΔΔG_f_^app^ = 5.1 kcal/mol) with respect to the wild-type (WT), most probably due to the loss of the stabilizing interaction between H298 and the aromatic ring of F295 which is not present upon the F to L substitution. In addition, both G395S and G395A variants, belonging to the loop connecting helix 8 and AF2, displayed a significantly smaller variation in folding free energy of the active form compared to the auto-repressed conformation. Such differences arise from the different local conformation of the two states; in the auto-repressed form the loop is tightly packed between AF2 and helix 8, while in the active form, the fusion of helices 7 and 8 (**Figure 1B**) pushes residue G395 outside of the protein core, thus rendering the residue significantly more solvent-exposed and therefore more tolerant towards any potential mutation.

The structural impact of the LBD nonsense mutations, resulting in the truncation of the protein at the level of Q244 and E400, was evaluated by running exhaustive all-atom 500 ns MD simulations (**Figure 2**). The analysis of the truncated structures highlighted that out of the total 9 α-helices constituting the LBD (**Figure 2A**), the E400* variant (**Figure 2B**) lacked only the 9^th^, C-terminal helix (residue 399-408), that is the AF2 helix (**Figure 2A**, cyan), and therefore retained the first 8 α-helices, whereas the Q244* variant retained only the first two (**Figure 2C**), namely helix 1 (residues 183-194; marked in yellow in **Figure 2A**) and helix 2 (residues 219-236; purple in **Figure 2A**). In terms of the structural evolution over the simulated timeframe, the Cα Root-Mean Square Deviation (RMSD) profile calculated with respect to the equilibrated structure (**Figure 2D**) suggested that the E400* truncation was overall slightly less prone to structural rearrangements over time compared to the WT form, with a ∼1 Å lower RMSD in the final 50 ns of the trajectory. On the contrary, the Q244* truncation displayed an abrupt increase in RMSD over the first 10 ns of simulation to values exceeding 6 Å, followed by a further increase to ∼ 9.5 Å during the rest of the trajectory (**Figure 2D**). This strongly points towards a significant decrease in the structural stability compatible with the loss of the largest part of the secondary structure elements due to truncation. The visual inspection of the trajectory of the Q244* variant highlighted a major rearrangement of the N-terminal helices 1 and 2 with respect to the WT, as shown by the different relative orientation of the helices, whose angle exhibited a significant increase from 140.5 ± 7.7 ° in the case of the WT to 157.1 ± 12.5 ° for the Q244* variant (**Figure 2C**). Concerning the E400* variant, no hints of a potential unfolding were observed, as shown by the preservation of the overall topology of the α-helices. However, a small but significant reduction in the amplitude of the angle between helices 1 and 2 and between helices 1 and 8 was observed in the E400* variant (140.5 ± 7.7 ° to 135.4 ± 7.7 ° and 33.5 ± 6.4 ° to 29.3 ± 7.7 °, respectively, **Figure 2B**), suggesting that the truncation allosterically affects the N-terminal region, as well as the proximal helix 8.

**Figure 2.**
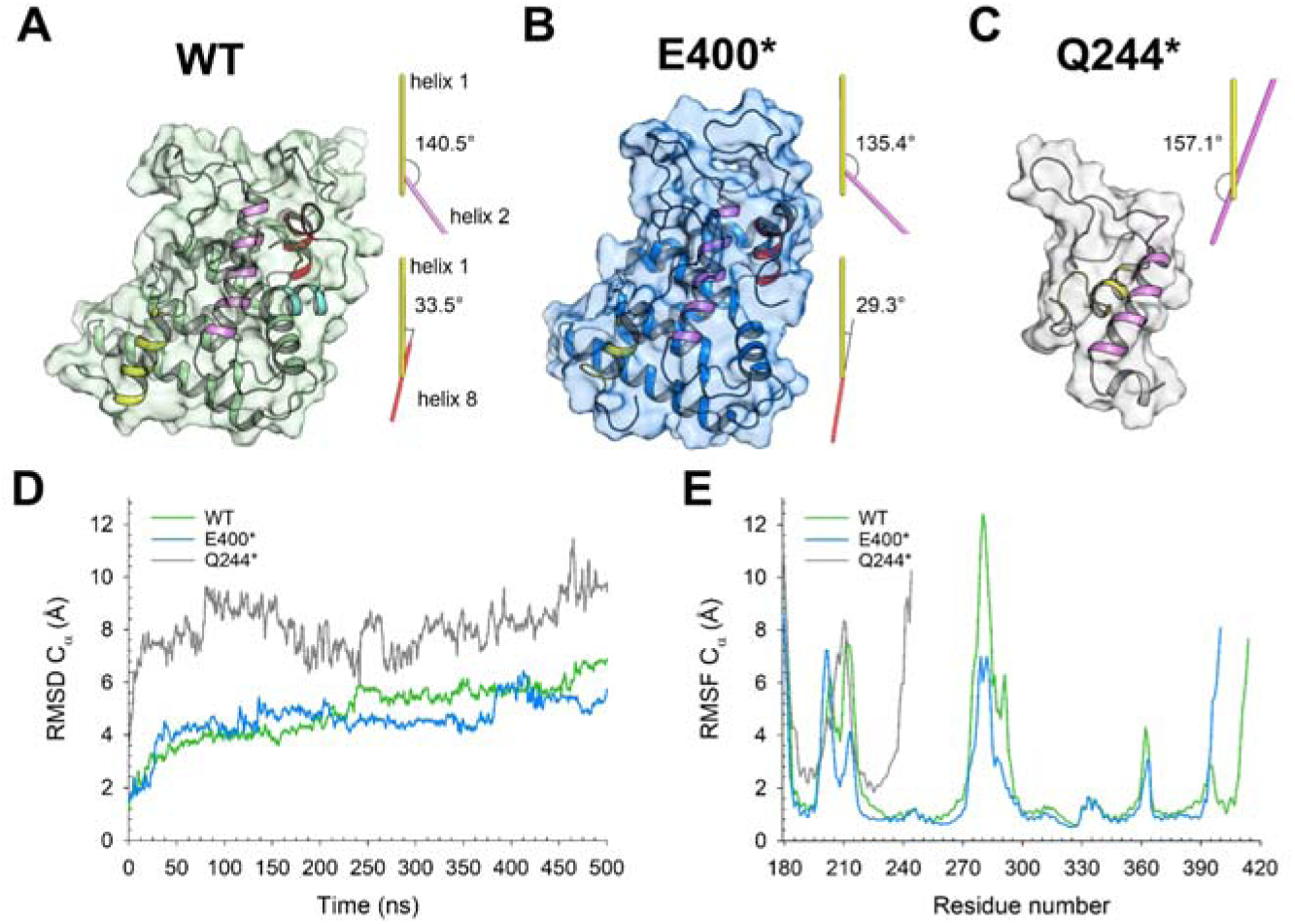
Three-dimensional structure of NR2F1 LBD. (**A**) WT (green), (**B**) E400* (blue), and (**C**) Q244* (grey) variants after 500ns MD simulations. Protein structure is shown as a cartoon with the molecular surface in transparency, helices 1 (residues 183-194), 2 (residues 219-236), 8 (382-394), and AF2 (residues 399-408) are colored in yellow, purple, red, and cyan, respectively. Insets show the schematic representation of the angles between helix 1 and helix 2 (yellow and purple) and between helix 1 and helix 8 (yellow and red) and their relative values. (**D**) Root-Mean Square Deviation of Cα atoms calculated over 500ns MD simulations with respect to the equilibrated structure of NR2F1 LBD WT (green), E400* (blue), and Q244* (grey). (**E**) Root-Mean Square Fluctuation of Cα atoms calculated over 500 ns MD simulations of NR2F1 LBD WT (green), E400* (blue), and Q244* (grey).

We also evaluated the flexibility of the backbone of the LBD by monitoring the Cα Root-Mean Square Fluctuation (RMSF), which represents the RMSD of each Cα (calculated with respect to their counterpart in the average structure) averaged over the number of frames in the MD trajectory (**Figure 2E**). Besides the Q244* truncation, which showed the largest plasticity throughout the entire structure, the E400* variant displayed less flexibility at the level of loops 195-218 and 270-294 compared to the WT, again indicating an allosteric effect due to the truncation. Interestingly, negligible differences were found in the flexibility of the LBD dimerization interface (residues 340-380), suggesting that the dimerization of the E400* variant may not be affected by the absence of the AF2 helix.

Taken together, our data suggest that both truncations are associated with BBSOAS through different molecular mechanisms. Since the Q244* variant preserves only the DBD, the eventual DNA-binding process is rendered independent of LBD regulation, whereas the E400* variant acquires a more compact conformation, which is however unable to switch from the auto-repressed to the active conformation, leaving the CRS available to any potential binding partner with no possible further regulation.

### LBD truncations and point mutations differently affect cell proliferation and survival

Having predicted the molecular effect of BBSOAS-associated LBD mutations by structural analyses, we further evaluated the impact of selected NR2F1 mutations at the cellular level to gain insight into the NR2F1 structure-function relationship (**Figure 3**). We focused on the Q244* and E400* truncations, and five distinct point mutations selected with respect to (i) their spatial locations, i.e., CRS (L252P), between CRS and DI (E318D), DI (G368D, L372P), and near AF2 (G395A) (**Figure 3A**), (ii) the diverse clinical symptoms with which they are associated (**Tables 1 and S1**), and (iii) their variable impact on isolated protein stability and affinity of both auto-repressed and active forms based on our computational analysis (**Tables S2 and S3**).

**Figure 3.**
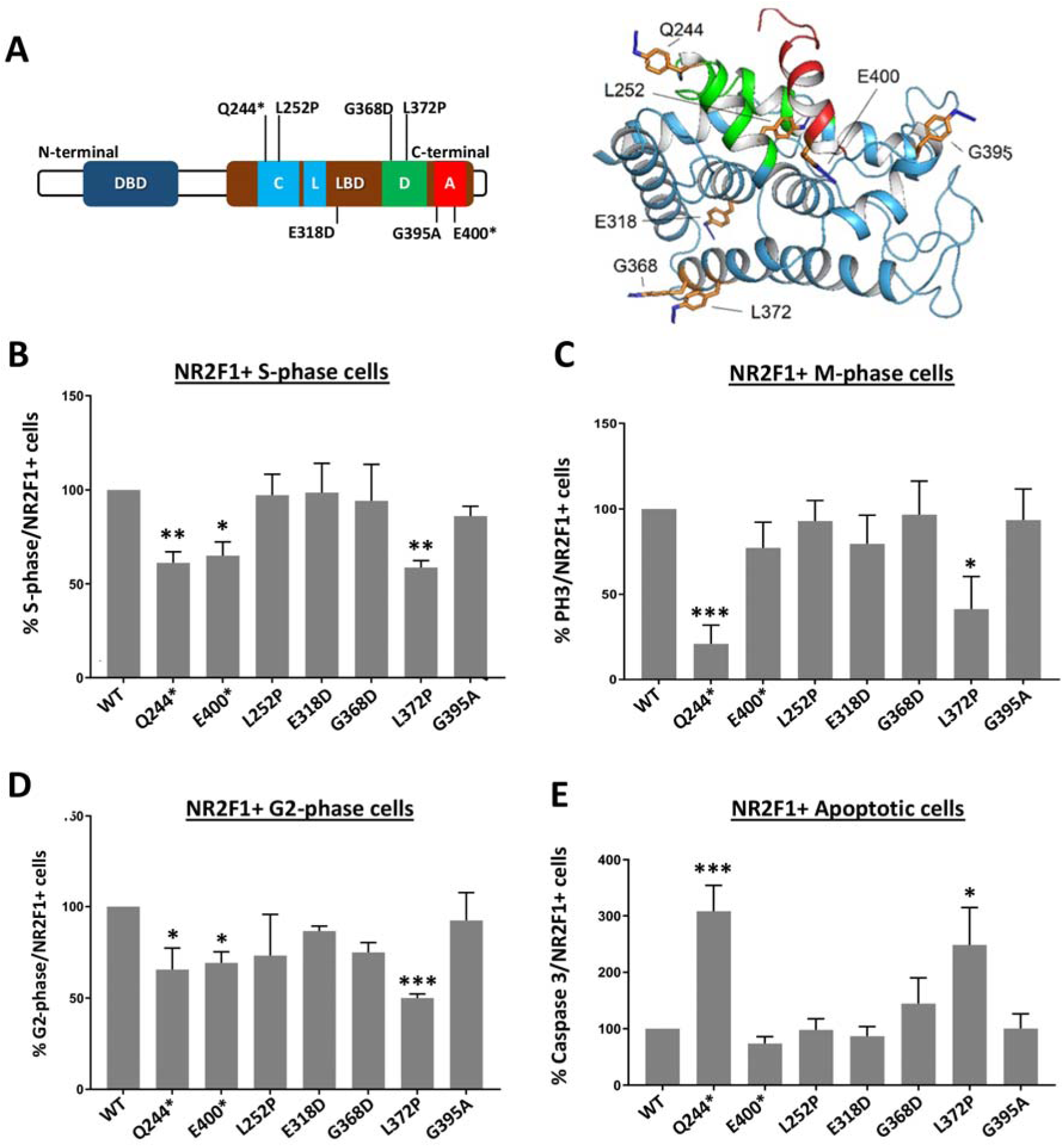
The impact of NR2F1 LBD mutations on cell physiology. (**A**) Left: Schematic diagram of mutation positions investigated (C, putative coactivator recognition site, L, putative ligand-binding site, D, dimerization surface, A, AF2), Right: Three-dimensional structure of the LBD shown as a cyan cartoon with the AF2 helix shown in red and the CRS in green. Amino acid positions investigated are shown as orange sticks with N atoms in blue and labeled according to the original residue present in that position. (**B-D**) Cell cycle analysis upon transient transfection of HEK293T cells with the different NR2F1 variants as indicated. Histograms showing the percentages of S-phase (**B**), M-phase (**C**) and G2-phase (**D**) cells in NR2F1-positive populations. (**E**) Flow cytometry analysis of NR2F1-positive cells staind for anti-cleaved-caspase 3 and undergoing apoptosis. FACS gatings are presented in **Figure S3**. Means ± SD of at least 3 independent experiments are shown (∗p<0.05, ∗∗p<0.01, ∗∗∗p<0.0005, one-way ANOVA).

To investigate the effect of these pathogenic NR2F1 mutations on cell physiology, we took an *in cellula* overexpression approach by transfecting the different NR2F1 variants in HEK293T cells, which expressed marginal levels of endogenous NR2F1 (not shown). First, by means of double labeling with propidium iodide and NR2F1, we quantified the fraction of NR2F1-positive cells entering different cell cycle phases by flow cytometry (**Figure 3B-D**; **Figure S3A, B**). Interestingly, HEK293T cells transfected with the Q244*, E400*, and L372P variants showed a significant decrease in S- and G2-phase entry when compared to WT NR2F1, whereas other variants only showed a trend towards reduction (**Figure 3B,D**), suggesting that distinct NR2F1 mutated forms can significantly inhibit cell cycle progression. By means of a phospho-Histone-3 (PH3) staining, we also assessed M-phase mitotic cells (**Figure S3C**). Upon transfecting the Q244* and L372P forms, we found a significant decrease in PH3-positive cells in the NR2F1-positive population (**Figure 3C**), indicating that cells carrying these variants fail to reach the M-phase and accomplish cell division. These findings suggest that some patient-specific LBD variants affected cell cycle progression, by slowing down to different extents the entry into the S-, G2- and/or M-phase.

We reasoned that a blockage in the cell cycle progression could be associated with a higher tendency to undergo cell apoptosis. Hence, we coupled the NR2F1 staining with cleaved Caspase-3 detection and quantified the percentage of apoptotic cells in the NR2F1-positive fraction (**Figure S3D**). Interestingly, the same two variants that affected the M-phase entry (the Q244* and the L372P) also significantly induced apoptosis (**Figure 3E**), indicating that a severe cell cycle disruption due to these two variants might ultimately result in apoptosis. Notably, the consequences of slowed cell cycle progression and increased apoptosis were readily visible in the cultured HEK293T cells 48 hours after transfection, with the Q244* variant inducing the most striking phenotype (**Figure S4**).

Together, these results demonstrate that specific NR2F1 truncations and point mutations had direct consequences on cell proliferation and survival and might suggest a toxic *gain-of-function* mechanism that could partially explain how specific NR2F1 variants impact human cell physiology.

### LBD truncations and point mutations can disturb the sub-cellular localization of NR2F1

Next, we assessed whether the presence of pathogenic mutations could affect the normal sub-cellular localization of NR2F1 in the nucleus. To this end, we co-immunostained NR2F1 in transfected HEK293T cells with DAPI (nuclear) and an anti-acetylated tubulin (cytoplasm/cytoskeleton) antibody and quantified the fraction of cells displaying NR2F1 either in the nucleus and/or cytoplasm (**Figure 4**). As expected from a transcriptional regulator, WT NR2F1 was mainly localized in the nucleus (**Figure 4A**), with approximately 80% of the cells showing this subcellular localization (**Figure 4B**). Strikingly, the vast majority (70%) of the Q244*-transfected cells clearly displayed NR2F1 cytoplasmic staining (**Figure 4A**, C), while only a minor fraction (25%) revealed a partial NR2F1 localization both in the nucleus and the cytoplasm (**Figure 4A**, D). Notably, the L252P point mutation and to a minor extent the E318D and E400* mutations showed intermediate phenotypes with a partial accumulation of NR2F1 in both cytoplasm and nucleus (**Figure 4A**, D). On the contrary, other NR2F1 variants (G368D, L372P, and G395A) resulted mainly localized in the nucleus, similar to the NR2F1 WT (**Figure 4A**, B), suggesting that the presence of these mutations does not affect subcellular distribution.

**Figure 4.**
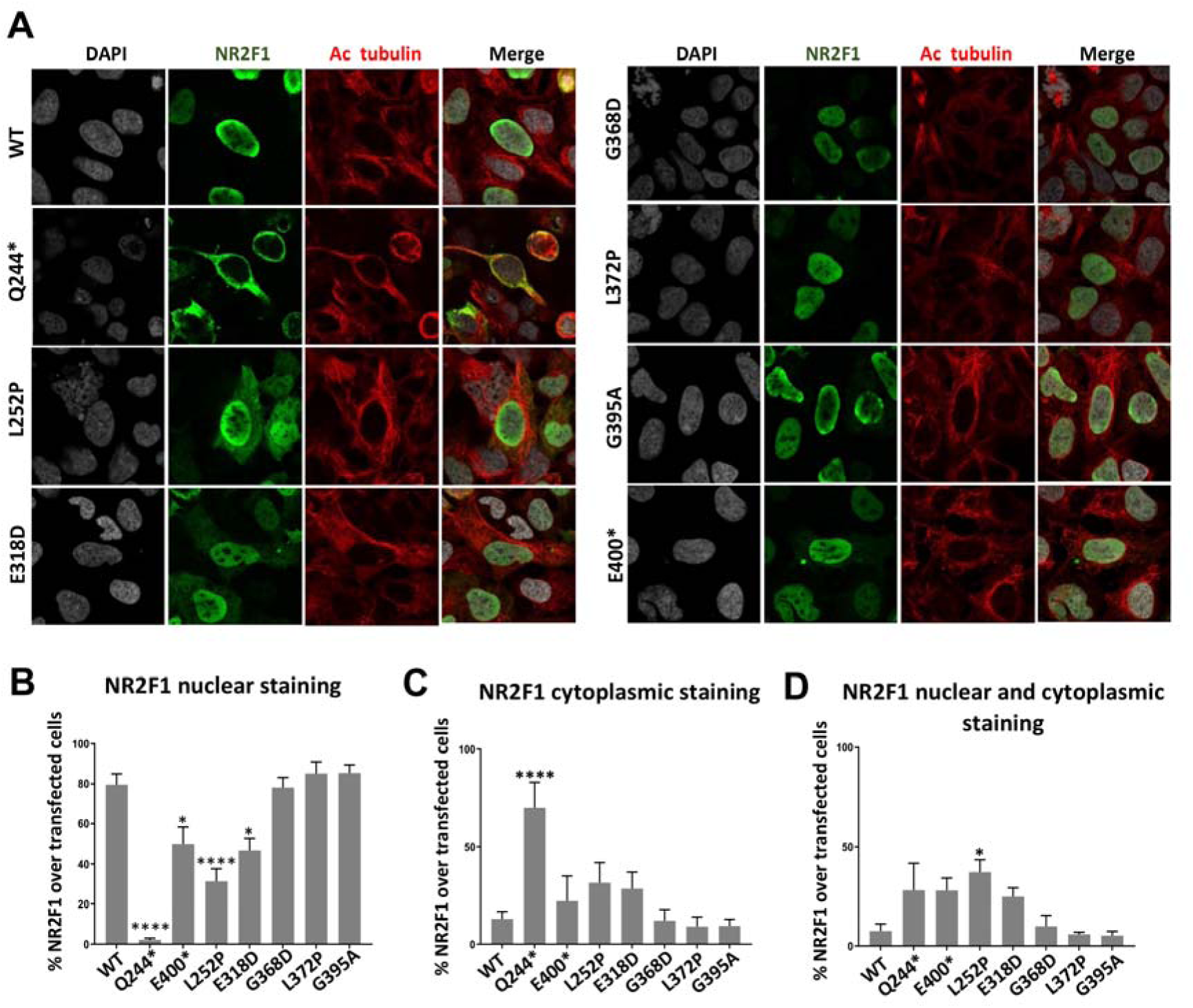
The impact of NR2F1 LBD mutations on NR2F1 localization. (**A**) NR2F1 (green) and acetylated tubulin (red) immunofluorescence staining on HEK293T cells transiently overexpressing NR2F1 (WT or mutants), as indicated on the side. (**B-D**) Histograms showing the quantification of the percentage of NR2F1 subcellular location in the nucleus (**B**), in the cytoplasm (**C**), and in both the nucleus and cytoplasm (**D**). Only NR2F1-positive cells were counted. Means ± SD of three independent experiments are shown (∗p<0.05, ∗∗∗∗p<0.0001, two-way ANOVA).

Taken together, these results indicate that only some LBD mutations/truncations affect NR2F1 nuclear localization. Notably, such abnormal intracellular distribution does not always mirror the presence of abnormal cell proliferation and survival. Indeed, our data suggest that proper nuclear localization alone is not indicative of proper NR2F1 functions since NR2F1 pathogenic variants can exert toxic effects both when accumulating in the cytoplasm (as for Q244*) as well as when normally localized in the nucleus (as for L372P).

### Impact of LBD mutations on the formation of NR2F1-protein complexes

It is generally accepted that most nuclear receptors undergo homo- or heterodimerization with other nuclear receptors before or upon DNA binding (Chandra *et al*, 2017; Lou *et al*, 2014) through the LBD domain (modeled in **Figure 1D**). Following the notion that NR2F1 binds DNA as a homo- or heterodimer (Park *et al*., 2003; Tsai & Tsai, 1997), any alteration of the dimerization process is expected to significantly interfere with the cellular function of NR2F1. In this line, the observation that the Q244* variant affected cell survival and proliferation more strongly than the E400* variant was thus not surprising, as the former lacks most of the protein interaction interfaces in the LBD, while the latter lacks only the AF2 helix.

To further assess the impact of NR2F1 pathogenic mutations on the capacity of forming dimers which may also affect the cellular function, we evaluated the effect of LBD mutations on the ability of NR2F1 to dimerize with itself, with NR2F2 and/or with RXRα (Evans & Mangelsdorf, 2014). To first investigate NR2F1 homodimerization, we co-expressed FLAG-tagged NR2F1 (FLAG-NR2F1-WT or mutants) and Myc-tagged WT NR2F1 (Myc-NR2F1) in HEK293T cells followed by co-immunoprecipitation (co-IP) with anti-FLAG antibody and immunoblotting. The detection of Myc-NR2F1 coimmunoprecipitated with FLAG-NR2F1 on the immunoblot would indicate a complex formation among the NR2F1 molecules (e.g., homodimers or oligomers). Surprisingly, we found that the co-IP of Myc-NR2F1 with FLAG-NR2F1-WT was barely detectable (**Figure 5A**), as with the FLAG-NR2F1 G368D and G395A variants. On the contrary, significant amounts of Myc-NR2F1 co-IP with the FLAG-NR2F1-L252P, E318D, and L372P variants were observed, even though expression levels of these variants were consistently lower than that of the WT. By performing densitometric analysis followed by normalizing the Myc-NR2F1 by FLAG-NR2F1 co-IP levels, we found that L252P, E318D, and L372P variants increased the complex formation by 12-fold, 20-fold, and 8-fold, respectively, when compared to WT NR2F1 (**Figure 5B**). Interestingly the expression of the G395A variant consistently resulted in a reduced NR2F1/NR2F1 complex formation although its expression level was always higher than that of the WT NR2F1, given the same transfection parameters.

**Figure 5.**
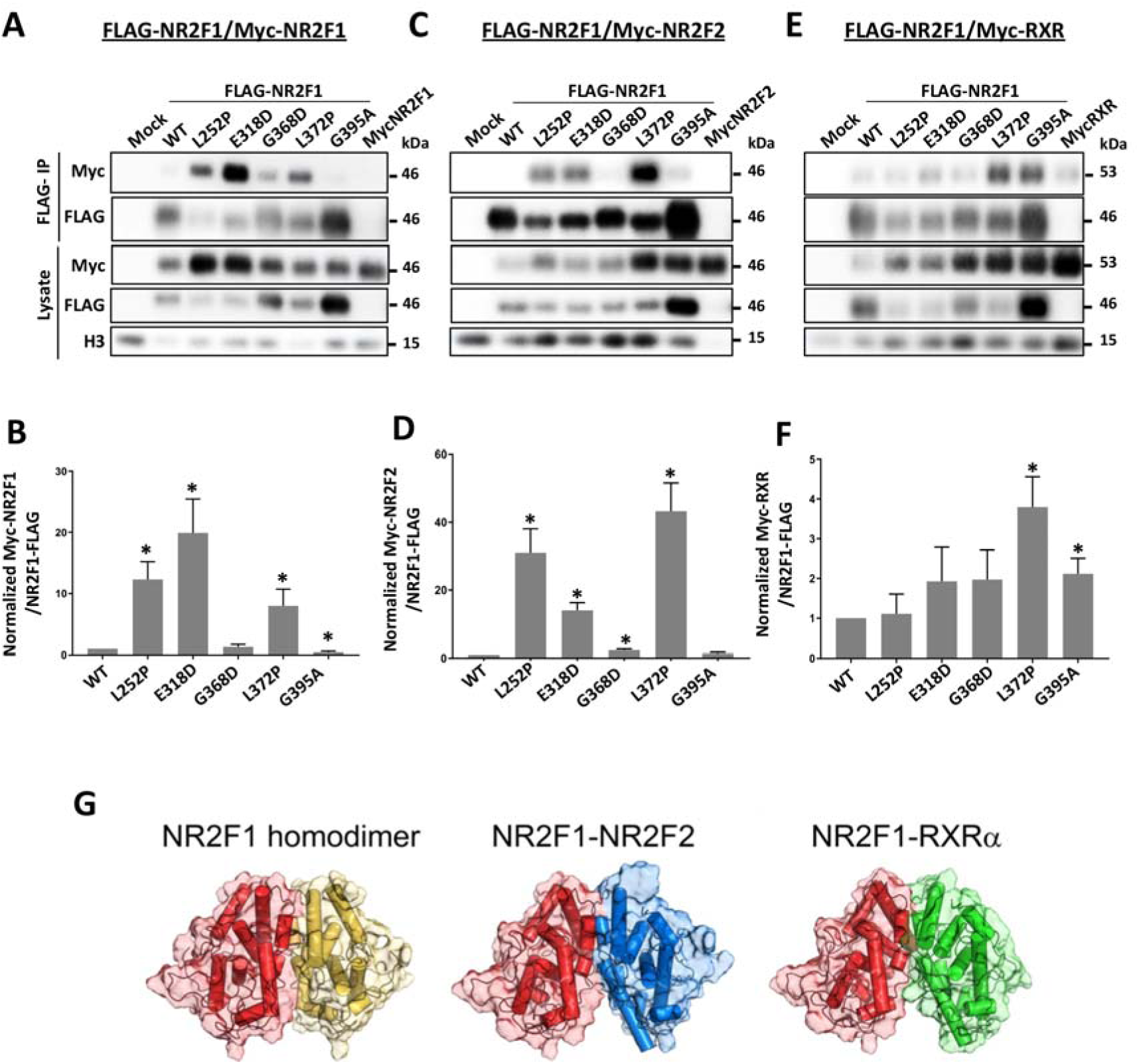
The impact of NR2F1 LBD mutations on its oligomerization in cells. (**A, C, E**) Immunoblots of lysates from transiently co-transfected HEK293T cells with NR2F1-FLAG (WT or mutants) and Myc-NR2F1 (**A**), Myc-NR2F2 (**C**), or Myc-RXRα (**E**). (**B,D,F**) Densitometric analysis of coIP samples from experiments as shown in A), C), and E) using ImageJ. Intensity normalization was done by first dividing the band intensity of each co-IP Myc-NR2F1 variant by the intensity of the corresponding FLAG-NR2F1 (Myc-NR2F1/FLAG-NR2F1). The resulting ratio of each variant was then normalized by the wild-type Myc-NR2F1/FLAG-NR2F1 ratio, such that the normalized wild-type Myc-NR2F1/FLAG-NR2F1 was equal to 1. Means ± SD of at least 3 independent experiments are shown (∗p<0.05, unpaired t-test, compared to NR2F1-WT). **G**) Results from docking simulations of NR2F1 LBD with itself (left), NR2F2 LBD (center) and RXRα LBD (right), proteins are represented as cartoon with cylindrical helices together with the respective molecular surface in transparency, NR2F1 LBD is shown in red and yellow, NR2F2 LBD in blue and RXRα LBD in green.

Then, we performed similar experiments to investigate the NR2F1 heterodimerization with its homolog NR2F2 and with the nuclear receptor, RXRα, previously described as potential NR2F1 dimerization partners (Cooney *et al*., 1992; Pinaire *et al*, 2000; Zhang & Dufau, 2001), by co-expressing FLAG-NR2F1 (WT or mutants) with Myc-tagged NR2F2 (Myc-NR2F2) or Myc-tagged RXRL (Myc-RXRα) (**Figure 5C-F**). As with the NR2F1 homodimerization experiments, we could not readily detect a WT NR2F1/2 interaction (**Figure 5C**). However, we observed a 31-fold, 14-fold, and 43-fold increase in the NR2F1/2 complex upon the expression of L252P, E318D, and L372P variants, respectively, according to densitometric analysis followed by normalizing Myc-NR2F2 by FLAG-NR2F1 co-IP (**Figure 5D**). Additionally, we also detected a small but significant increase (∼2-fold) in NR2F1/2 co-IP upon the expression of NR2F1-G368D while no increase was observed for the NR2F1 G395A variant. Consistently, while we did not observe significant RXRα interaction with WT NR2F1 nor NR2F1-G368D above the background level, we observed an increase in RXRα interaction with NR2F1-L372P (∼3-fold) (**Figure 5E, F**). However, unlike NR2F1/NR2F1 or NR2F1/2 interactions, the NR2F1 L252P and E318D variants did not significantly enhance NR2F1/RXRα interaction. Notably, while the expression of the G395A variant reduced the NR2F1 homo-oligomerization by about ∼60% (**Figure 5B**), inversely, it significantly increased the NR2F1/RXRα interaction by approximately 2-fold (**Figure 5F**).

Overall, our data in HEK293T cells suggest that NR2F1 homo- and heterodimerization may not always be a prevalent event and that some NR2F1 pathogenic variants may instead promote the protein propensity to form homo-/heterodimers and/or higher order NR2F1 complexes. Furthermore, depending on the identity of the dimerization/oligomerization partners, different variants may exert varying degrees of impact on their interactions.

### *In silico* NR2F1 protein complexes modeling

To further interpret the above-described experimental results, we predicted the quaternary structure of NR2F1 variants *in silico*. Since no experimental structure of NR2F1 dimers was available, we used PIPER (Kozakov et al., 2006) to run unbiased protein-protein docking simulations. Simulations revealed a dimeric assembly essentially in line with the conserved dimerization interfaces between other proteins belonging to the same nuclear hormone receptor superfamily, namely NR2F2, RXRα (subfamily NR2), RARβ, and LXRβ (subfamily NR1). The similarity concerned both the sequence (**Figure S1**) and the physicochemical nature of the interactions between monomers (**Figure S2**). Based on the structural information provided by the experimental structures of the NR2F2 homodimer (Kruse *et al*., 2008), RXRα homodimer (Egea *et al*, 2002), RXRα-RARβ (Chandra *et al*., 2017), and RXRα-LXRβ (Lou *et al*., 2014) heterodimers (**Figure S2**), we predicted a reliable three-dimensional structure of NR2F1 homodimer and the heterodimeric complexes of NR2F1 with NR2F2 and RXRα (**Figure 5G**), both in the auto-repressed and active forms. To probe the robustness of predicted NR2F1 LBD homodimers in both the auto-repressed and the active forms, rigid-body docking simulations were performed using another docking algorithm, namely ZDOCK 2.3 (Chen *et al*, 2003), whose score (ZDOCK-score) was demonstrated to correlate with the experimental binding affinity upon an exhaustive sampling of the roto-translational space and under the assumption of rigid body-like binding (Dell’Orco *et al*, 2007). We were thus able to identify for LBD in both states, 23 poses resembling the docked complex obtained with PIPER (native-like poses, Cα-RMSD < 1 Å, **Table S4**). The active form presented higher-ranked best solutions compared to the auto-repressed form in all 4 docking runs (**Table S4**), which reflected the higher average ZDOCK-score of the native-like solutions of the active form (58.06 ± 6.63 vs 49.31 ± 3.97, **Table S4**). No direct estimate of the binding free-energy was possible in this case, since the original affinity-ZDOCK-score correlation model was obtained with protein-protein interfaces significantly smaller than those of NR2F1 homodimers (Dell’Orco et al., 2007). Nevertheless, the ZDOCK-score values were higher than those obtained with the same protocol for another small globular protein (Guanylate Cyclase Activating Protein 1), whose dimeric architecture was validated by a combination of analytical Size Exclusion Chromatography coupled to Multi-Angle Light Scattering, Small-Angle X-Ray Scattering, and Microscale Thermophoresis (Boni *et al*, 2020). Thus, independent and unbiased docking simulations suggest for the NR2F1 LBD a dimerization process substantially in line with that of other nuclear receptors in solution, both structurally and in terms of affinity (**Figure 5G**).

Based on the predicted structural organization of NR2F1 homo- and heterodimers, we then calculated the effects of BBSOAS-variants relative to the WT on quaternary complex structures, in terms of folding stability (ΔΔG_f_^app^) and binding affinity (ΔΔG_b_^app^). Among all tested variants, with the exception of T200R, the free energy of folding ΔΔG_f_^app^ of the homo/heterodimers with respect to the WT in both the auto-repressed and active conformation was increased, regardless of the dimerization partner (see **Video S3** for results calculated on NR2F1 homodimer in heterozygosis as an example). This suggests a destabilization of the LBD dimeric complexes in the presence of BBSOAS-associated variants, with the largest effect being displayed by the NR2F1 homodimer with mutations in a potential homozygosis genetic condition (**Tables S2 and S3**).

Interestingly, a generally lower effect on the dimerization affinity was predicted for BBSOAS-associated variants, as only R366C, G368D and L372P displayed significantly diminished affinity (i.e. higher ΔΔG_b_^app^ values) for each of the tested complexes in both the active and the auto-repressed conformations, although with numerical differences among the partner receptors. This is not surprising, as all three residues belong to helix 7, which constitutes the dimerization interface together with helices 6 and 8. Moreover, such mutations displayed an overall more prominent decrease in affinity in the case of the active form (**Table S3**) compared to the auto-repressed one (**Table S2**), most likely due to the increased interaction surface. Interestingly, the E342K mutation turned out to be detrimental only for the active form, where the larger dimerization interface brings residue 342 in close contact with the positively charged residues K308, K369, and R373 of the other subunit. The E342K variant would therefore replace a negative charge with a positive one, thus accounting for electrostatic repulsion to be responsible for decreasing dimer affinity. All other tested variants showed a negligible effect on dimer affinity, with ΔΔG_b_ ^app^ values never exceeding ± 1 kcal/mol (**Tables S2 and S3**).

Taken together, our results suggest that the predicted destabilizing effects of the point mutations on the isolated forms of NR2F1 correlate generally well with the propensity of NR2F1 to form dimers and/or oligomers in cells, although dimeric interfaces that differ from the canonical interface experimentally observed in crystal structures of other nuclear receptors may form under specific conditions.

### Participation of the amino acid positions affected by BBSOAS mutations in the dimerization as revealed by GCE-enabled photocrosslinking in living cells

Several putative protein interaction sites in the LBD are expected to allow NR2F1 to dynamically work in concert with various protein partners depending on the cellular stages and context. However, our conventional co-IP approach could not easily detect homo- and/or heterodimerization of predicted partners with the WT NR2F1 form despite the *in silico* prediction of the event (**Figure 5**), indicating the limitations of the approach to stably capture very transient interactions in living cells.

To overcome these experimental constraints and more precisely study protein interactions at selected BBSOAS mutation positions in a more physiologically relevant manner, we applied the genetic code expansion (GCE) technology to co-translationally and site-specifically incorporate a small photo-crosslinking amino acid, *p*-azidophenylalanine (AzF) into the LBD of NR2F1 (**Figure 6A**). This was achieved by co-transfecting HEK293T cells with NR2F1 isoforms (*NR2F1am*), each carrying an amber codon (*TAG*) at a desired position, along with an amber suppressor tRNA derived from *B. stearothermophylus* tRNA^Tyr^ (Bst-Yam) and an enhanced aminoacyl tRNA synthetase (E2AziRS) in the presence of AzF (Seidel *et al*, 2017). A subsequent UV irradiation allowed a photo-activated covalent crosslinking between AzF in the LBD and its interactors within the radius of reach of AzF (∼9 Å from CL) within a living cell (Seidel *et al*., 2017). We hypothesized that variable pathogenic effects of the BBSOAS mutations in different LBD protein-binding sites may be attributed to their interferences with different protein-protein interactions. Therefore, we focused our investigations on the same mutation locations at the protein interaction sites where we investigated the cellular effects (**Figures 3-4**).

**Figure 6.**
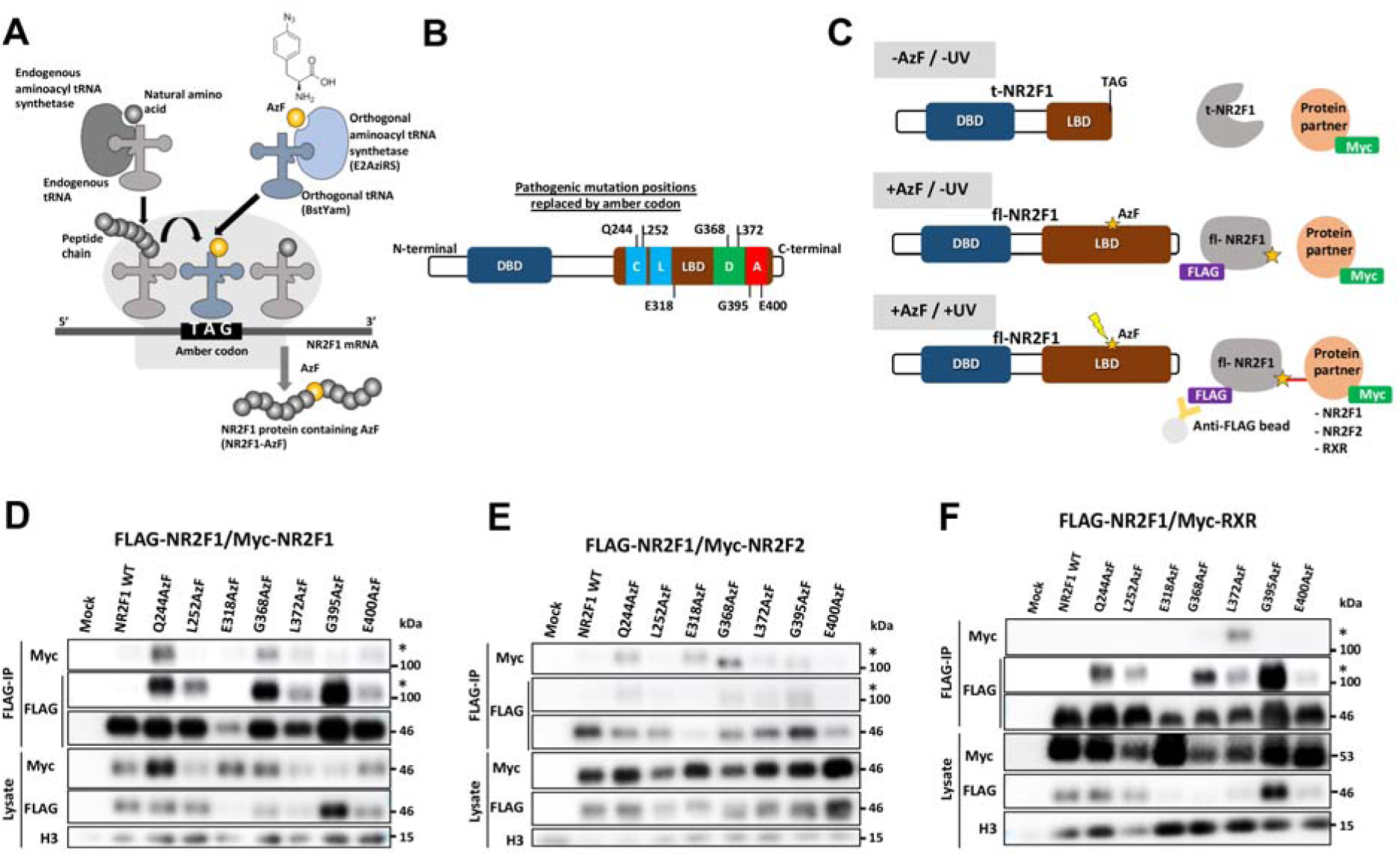
Covalent and site-specific NR2F1 dimeric partner capturing via GCE. (**A**) Principles of site-specific incorporation of photocrosslinking amino acid by amber codon suppression. Endogenous aminoacyl tRNA synthetase charges the endogenous tRNA with its cognate amino acid. Aminoacyl-tRNA enters the ribosome and adds the amino acid to the corresponding codon. For the AzF incorporation in NR2F1, the orthogonal aminoacyl tRNA synthetase (E2AziRS) catalyzes the aminoacylation between AzF and the orthogonal tRNA (BstYam). The AzF-charged tRNA enters the ribosome and incorporates the AzF in response to the designated amber codon. The translation continues and produces full-length, AzF-containing NR2F1 protein. (**B**) Schematic diagram of pathogenic mutation positions replaced by amber codon for AzF incorporation. (**C**) Diagram showing UV-induced, site-specific crosslinking via AzF. NR2F1 is produced as a truncated form (t-NR2F1) in the absence of AzF due to the designated amber codon. In the presence of AzF, the amber codon suppression allows the incorporation of AzF at the selected amber codon position and the NR2F1 protein carrying AzF is translated into the full-length FLAG-tagged form (fl-NR2F1). Upon UV irradiation (365 nm), AzF forms a covalent bond between FLAG-NR2F1 and Myc-tagged putative partners (NR2F1, NR2F2, RXRα). The proteins in the crosslinked complex can be co-IP with NR2F1 by using anti-FLAG antibody-conjugated beads and detected in a higher molecular weight complex by immunoblot. (**D-F**) Immunoblots showing site-specific capturing of homodimer (FLAG-NR2F1/Myc-NR2F1 (**D**), heterodimers (FLAG-NR2F1/Myc-NR2F2 (**E**), and FLAG-NR2F1/Myc-RXRα (**F**). The expected position of the covalently bound dimers is denoted by an asterisk.

To this end, we generated a series of C-terminal FLAG-tagged full-length NR2F1 variants where the amino acid positions of mutations Q244*, L252P, G318D, G368D, L372P, G395A, and E400* were replaced by AzF in response to the amber codon (TAG) (**Figure 6B**). The incorporation of AzF was successful with minimal read-through in the absence of AzF, and NR2F1-interacting proteins could be covalently captured by photocrosslinking *via* AzF (**Figure S5**). Then, we compared the ability of different AzF-NR2F1 variants to capture the elusive dimerization *in cellula* by co-expressing FLAG-AzF-NR2F1 with Myc-NR2F1, Myc-NR2F2, or Myc-RXRα in HEK293T and then performing GCE-enabled photocrosslinking followed by FLAG-NR2F1 co-IP (strategy in **Figure 6C**). The results showed that the NR2F1 homodimerization and NR2F1/2 heterodimerization were most efficiently captured by Q244AzF and G368AzF variants, whereas the NR2F1/RXRα heterodimerization was most efficiently captured by the L372AzF variant (**Figure 6D-F**). Although some NR2F1/RXRα heterodimerization could be captured by the Q244AzF variant, this was less efficient (not shown). To further probe the conformational impact of pathogenic mutations on the dimer formation while avoiding possible functional interference by the photocrosslinker incorporation, we chose to place the photocrosslinker at the residue 244 (Q244AzF), since this residue was not predicted as a critical residue for dimerization (**Figures 1D**, 5G) but close enough to allow the capture of dimerization partners of WT NR2F1 without perturbing the cellular function of the protein (**Figure S6**).

To investigate the effects of point mutations L252P, E318D, G368D, L372P, and G395A on the dimer formation using the photocrosslinking *via* the Q244AzF, we co-expressed full-length FLAG-NR2F1 or FLAG-Q244AzF variants (with or without L252P, E318D, G368D, L372P, or G395A pathogenic mutations) with Myc-tagged NR2F1, NR2F2, or RXRα, and the *E2AziRS*/Bst-Yam pair in the presence of AzF (**Figure 7**). After photocrosslinking with UV, we performed co-IP of the cell lysates with the anti-FLAG antibody and immunoblotting. As expected, the photocrosslinking *via* the Q244AzF successfully captured the elusive NR2F1 homodimer and heterodimers with NR2F2 and RXRα, as indicated by an enrichment of the FLAG-Q244AzF-NR2F1/Myc-NR2F1, NR2F2, or RXRα complex at the molecular weight corresponding to the NR2F1 homodimer or heterodimer (**Figure 7A-C**). Notably, the presence of L252P, E318D, G368D, and L372P mutations clearly led to a reduction in NR2F1 protein expression. Nevertheless, these variants were detectable after immunoprecipitation. Densitometric analysis followed by normalizing Myc-NR2F2 by FLAG-NR2F1 co-IP indicated that the G368D mutation significantly reduced the ability of NR2F1 to form the homodimer (**Figure 7A**) as well as the heterodimer with NR2F2 and RXRα (**Figure 7 B-C**). This result confirmed the *in silico* prediction that this mutation exerted the strongest detrimental effect on dimer stability and affinity (**Tables S2 - 3**). Moreover, we did not observe a statistically significant effect of the L252P, E318D, and L372P mutations on the homo- and heterodimer formation even though the conventional NR2F1 co-IP showed that these mutations favored complex formation (**Figure 5A-C**). These observations suggest that the structural destabilizing effect of these pathogenic mutations not only led to the reduction of the NR2F1 protein expression but also increased the propensity of NR2F1 to form homo- and hetero-dimeric/oligomeric complexes that are not the dimeric complexes that could be captured by photocrosslinking at the Q244 position of WT NR2F1. Interestingly, of all the mutations tested, the G395A was the only mutation that caused a strong increase in NR2F1 protein expression levels. Although this mutation did not significantly affect the NR2F1 homodimerization or heterodimerization with RXRα (**Figure 7A**, C), it significantly reduced the NR2F1/2 heterodimerization (**Figure 7B**). Overall, the covalent and site-specific protein-protein interaction captured by GCE allowed us to discern the effects of different pathogenic mutations on distinct NR2F1 dimeric pairs in living cells.

**Figure 7.**
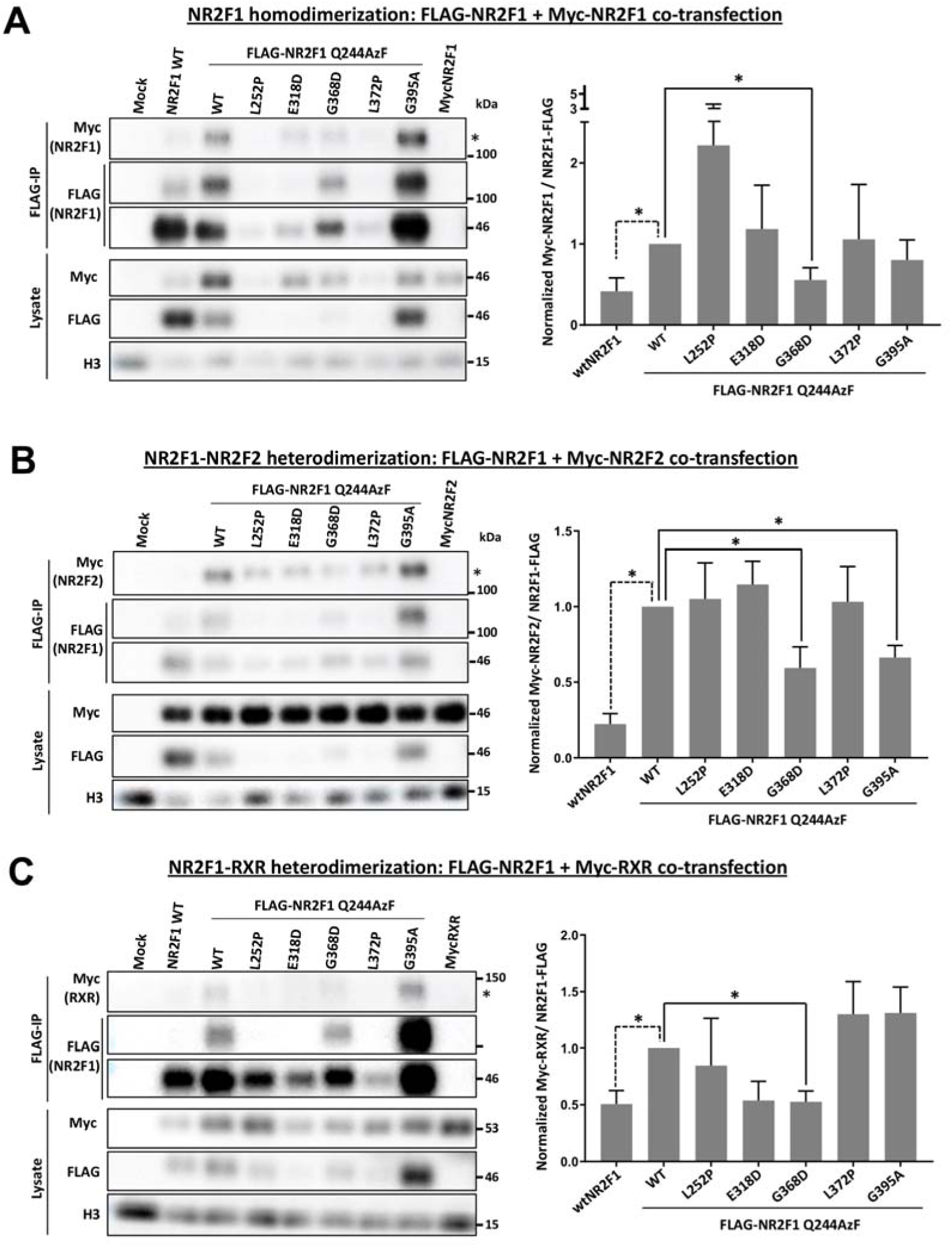
The impact of NR2F1 LBD mutations on its dimerization in living cells. (**A-C**) Panels showing immunoblots of the immunoprecipitated samples and whole lysates with anti-FLAG antibody. The pIRE4-Azi plasmid containing E2AziRS and BstYam, pcDNA3.1 NR2F1Q244*-FLAG without or with pathogenic mutations, and the plasmid carrying the gene of each putative protein partner (Myc-NR2F1, Myc-NR2F2, or Myc-RXRα) were transiently cotransfected in the HEK293T cells. Left: Immunoblots show the dimerization between FLAG-NR2F1 and its putative partners Myc-NR2F1 (**A**), Myc-NR2F2 (**B**), and Myc-RXRα (**C**). The asterisk indicates the approximated position of the dimer’s molecular weight. Right: Quantitative analysis of the impact of the pathogenic mutations on the dimeric binding between NR2F1 and NR2F1 (**A**), NR2F1 and NR2F2 (**B**), and NR2F1 and RXRα (**C**). The densitometric analysis and normalization were performed as described in **Figure 5**. Means ± SD of three independent experiments are shown (∗p<0.05, unpaired t-test, compared to NR2F1-Q244AzF).

### CRABP2 as a novel partner of NR2F1

Our data on the abnormal cellular localization of some NR2F1 variants suggest that the nuclear transport of NR2F1 might require a co-factor binding, which might be disrupted due to the presence of specific LBD mutations (**Figure 4**). In light of its sequence similarity with RXRα and RARb (**Figure S1**) and our results showing that NR2F1 could indeed heterodimerize with RXRα in living human cells (**Figure 7**), we hypothesized that NR2F1 might be directly or indirectly involved in retinoic acid (RA)-induced regulation, as previously suggested (reviewed in (Alfano *et al*., 2013)). The cellular retinoic acid-binding protein 2 (CRABP2) is responsible for the transport of RA to the nucleus and its delivery to other nuclear retinoic acid receptors, such as RXR and RAR (Sessler & Noy, 2005), and could represent a potential NR2F1 partner in this process. However, the dynamic shuttling of CRABP2 between the cytoplasm and the nucleus presents a challenge in detecting its transient interaction with nuclear receptors in living cells. By taking advantage of the GCE-enabled photocrosslinking *via* the Q244AzF-NR2F1 followed by endogenous CRABP2 co-IP and immunoblotting, we explored the potential interaction between NR2F1 and CRABP2 (**Figure 8**). We found that indeed CRABP2 could be covalently captured *via* photocrosslinking by Q244AzF-NR2F1, revealing the *in cellula* direct interaction between the two molecules through the CRS of NR2F1 (**Figure 8A**). Using the same FLAG-tagged NR2F1 mutations described above, we further examined the impact of pathogenic mutations on CRABP2/NR2F1 interaction. Upon overexpression of the G368D and G395A variants, we detected a marked increase in the NR2F1/CRABP2 complex. On the other hand, given the same experimental parameters, the opposite effect was observed for the L252P and L372P variants. The expression of the E318D variant, however, did not affect the extent of NR2F1/CRABP2 complex formation (**Figure 8B**). Importantly, in order to support the physiological significance of this interaction, we performed an immunofluorescence of NR2F1 and CRABP2 in embryonic E13.5 mouse forebrain and developing eye, two anatomical sites known to be affected in *Nr2f1* mutant mice and perturbed in BBSOAS patients (Bertacchi *et al*., 2022; Tocco *et al*., 2021). Strikingly, a clear NR2F1 and CRABP2 protein co-expression was observed in cells of the medial-posterior cortex from which the hippocampus will form, and of the developing neural retina (**Figure 8C-D**), supporting a physiological interaction between NR2F1 and CRABP2 during mouse development.

**Figure 8.**
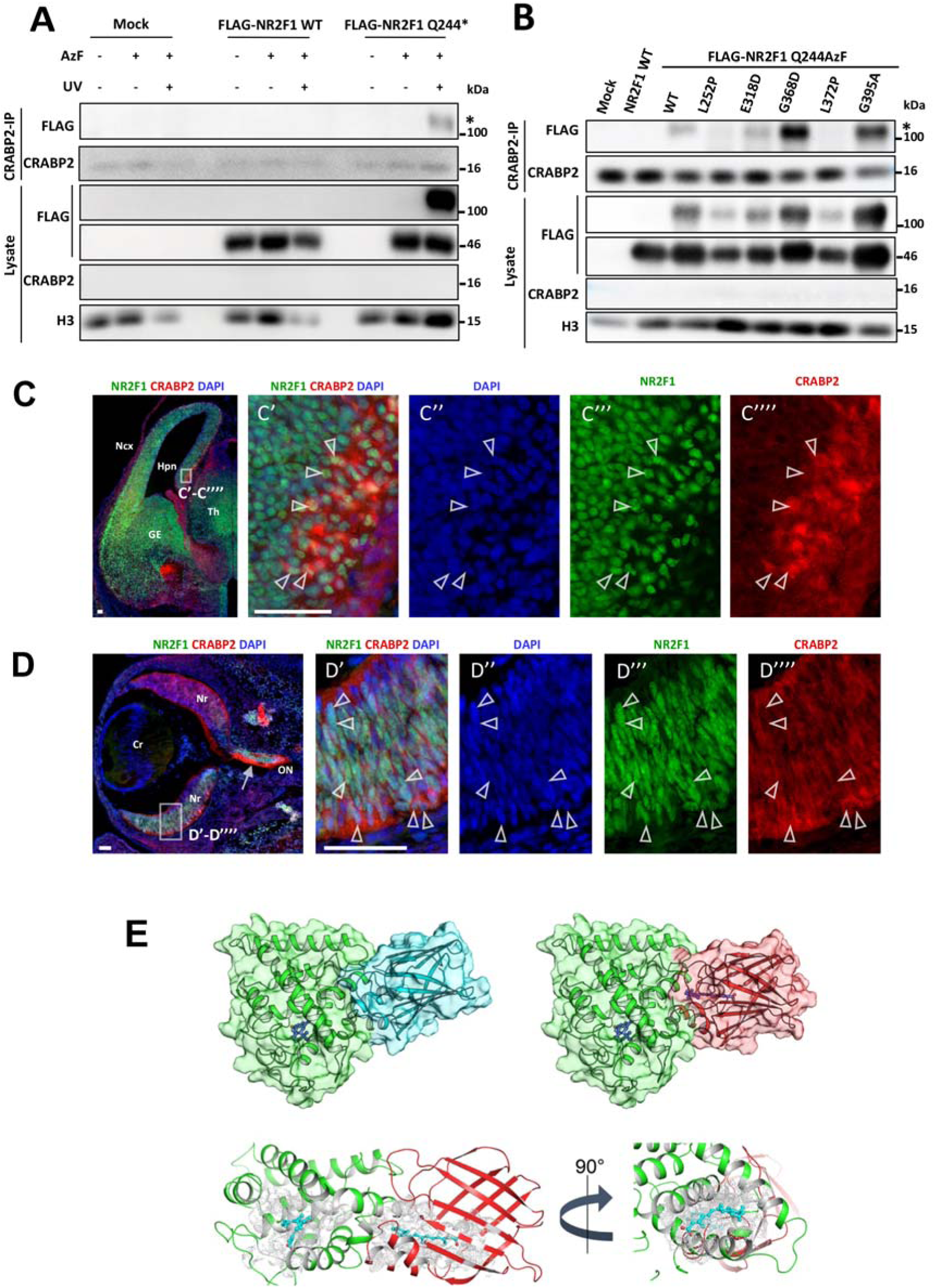
Direct interaction between NR2F1 and CRABP2. (**A)** HEK293T cells were transiently co-transfected with plasmids containing E2AziRS, BstYam, and FLAG-NR2F1-Q244*, subjected to CRABP2 coIP with anti-CRABP2 antibody, and immunoblotted with the indicated antibodies. The asterisk indicates the NR2F1/CRABP2 complex. (**B)** FLAG-NR2F1Q244* without or with pathogenic mutations was transiently co-transfected alongside E2AziRS and BstYam. Asterisks indicate the NR2F1/CRABP2 complex. **C)** NR2F1 (green) and CRABP2 (red) immunostaining of a WT mouse forebrain at E13.5. NR2F1 and CRABP2 co-expression is observed in cells of the hippocampal neuro-epithelium (Hpn) (**C’-C’’’’**). Arrowheads point to representative double-positive cells. Scalebar = 50µm. GE: ganglionic eminence; Ncx: neocortex; Th: thalamus. **D)** NR2F1 (green) and CRABP2 (red) immunostaining of a WT mouse neural retina (Nr) at E13.5. The white arrow in low magnification image points to high CRABP2 expression in optic nerve (ON) fibers, while arrowheads in high magnification images (D’-D’’’’) point to representative NR2F1/CRABP2 double-positive cells in the ventral Nr. Scalebar = 50µm. Cr: crystal lens. (**E)** Results from docking simulations of active NR2F1 LBD (bound to 9-cis retinoic acid) with apo- and holo-(bound to retinoic acid) CRABP2. Proteins are represented as cartoons together with the respective molecular surface in transparency, NR2F1 LBD is shown in green, apo CRABP2 in cyan, holo CRABP2 in red; retinoic acid and 9-cis retinoic acid are represented in sticks with C atoms in dark blue and O atoms in red. Structural detail of the hydrophobic pockets surrounding the ligands in the active NR2F1 LBD-holo CRABP2 complex. Protein structure is represented in a cartoon with active NR2F1 colored in green and holo CRABP2 in red, ligands are represented as sticks with C atoms in cyan and O atoms in red, the Van der Waals occupancy of each atom belonging to the hydrophobic pockets of either protein is represented as white dots. 90° counterclockwise view of the hydrophobic tunnel potentially allowing the transport of retinoic acid from CRABP2 to NR2F1 LBD.

Having discovered the interaction between NR2F1 and CRABP2 in cells and their co-expression in the mouse brain and eye, we further performed unbiased molecular docking simulations to identify potential interaction interfaces between CRABP2 and NR2F1. We compared the predicted binding interfaces of apo and holo CRABP2 (Vaezeslami *et al*, 2006) with NR2F1 and found that high-scored docking poses substantially shared the binding mode, suggesting that the interaction between NR2F1 and CRABP2 may occur regardless of the loading state of the RA transporter and without major conformational changes in either protein (**Figure 8E**). Moreover, the analysis of the hydrophobic pockets surrounding RA in NR2F1 and CRABP2 suggested that such pockets are aligned in the docked complex (**Figure 8E**), thus allowing RA to be directly channeled from CRABP2 to NR2F1 similarly to what occurs in the RA transport to RAR (Budhu & Noy, 2002). Interestingly, when the supramolecular NR2F1/CRABP2 assembly obtained by docking simulations was used to predict the effects of the experimentally tested point mutations on the stability of the protein complex, *in silico* results were in line with those obtained by GCE-enabled photocrosslinking, with a high destabilization predicted for the L252P and L372P variants (ΔΔG_f_^app^ =57.99 kcal/mol and 57.00 kcal/mol, respectively), a stabilizing effect for the G368D variant (ΔΔG_f_^app^ =-4.23 kcal/mol) and a negligible effect for the G395A variant (ΔΔG_f_^app^ =1.55 kcal/mol). This suggests that the interaction between NR2F1 and CRABP2 could be essential to create a functional RA channeling route similar to that observed in the complex with RAR, and that BBSOAS-associated variants may severely perturb the transport of RA. Taken together, these data support the physiological and pathological significance of the newly discovered interaction between the CRS of NR2F1 and CRABP2 in human cells and mouse embryos.

### Pathogenic mutations, protein interactions, functions, and clinical manifestation of BBSOAS

Drawing from the information we unveiled by structural and *in cellula* analyses regarding the effects of the pathogenic mutations, we looked for possible connections between the different mutations in NR2F1 LBD and the variation in the clinical manifestation of BBSOAS patients. While a definitive correlation could not be drawn due to the lack of sufficient clinical data, there are nevertheless some patterns that emerge (**Table 2**). While all the pathogenic mutations examined in the present study are associated with intellectual disability (ID) and most with developmental delay (DD), we can perceive that the additional set of symptoms that include visual impairment (VI), optic atrophy (OA), and optic disc/nerve (OD/ON) anomalies may be more associated with point mutations that strongly destabilized the isolated monomeric structure, reduced NR2F1 protein expression, and promote stable dimers/oligomers. These mutations (L252P, L372P, E318D) are those that exhibited perturbations in cellular functions investigated in this study, such as nuclear localization defect, cell cycle inhibition, and/or apoptosis. On the contrary, the additional set of symptoms that include epilepsy or seizure, autistic spectrum disorder or ASD-like symptoms, and motor delay (MD) may be more associated with point mutations that did not exhibit significant perturbations in the cellular functions investigated but reduced NR2F1 homodimer and/or NR2F1/2 or NR2F1-RXRα heterodimer (E318D, G368D, G395A) and may interfere with other cellular functions not covered here. The co-expression of NR2F1 and CRABP2 in the ventral retina and in other brain regions along with the observation that L252P and L372P mutations (which strongly inhibited CRABP2/NR2F1 binding) are associated with VI, OA, and OD/ON anomalies (while the G368D and G395A mutations are not) infer that the CRABP2-mediated NR2F1 functions may be one of the factors contributing to the visual symptoms of BBSOAS. Future investigation into the role of CRABP2-mediated NR2F1 is necessary to confirm this hypothesis.

**Table 2.**
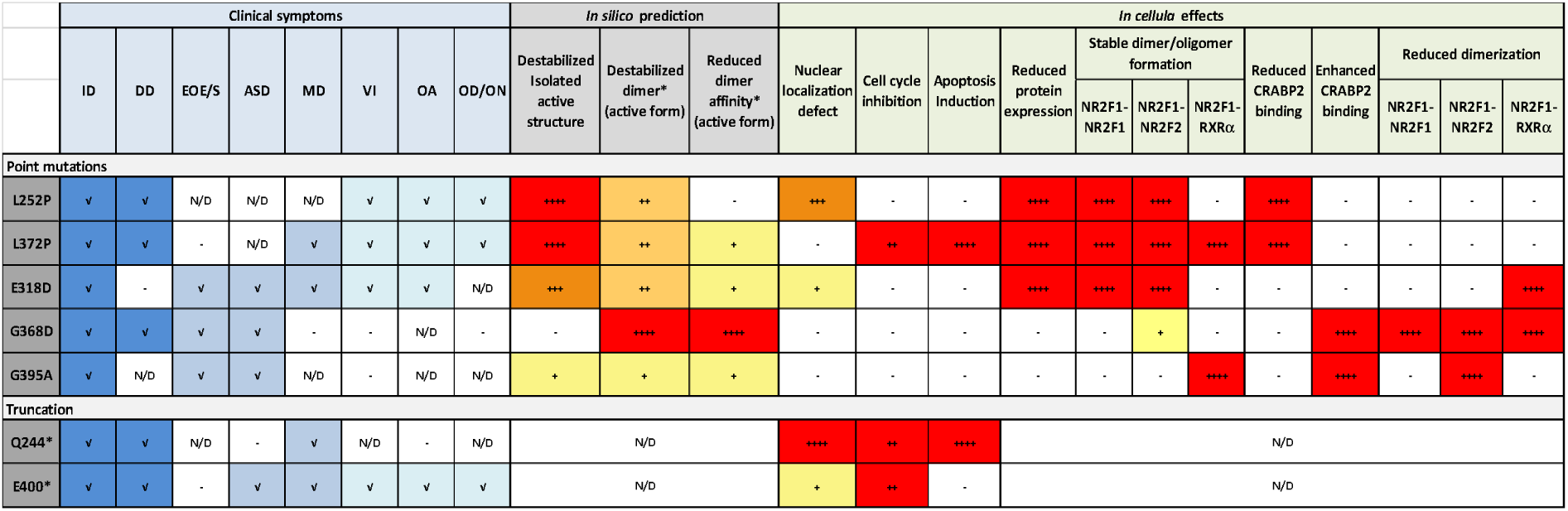
Summary of selected NR2F1 pathogenic mutations based on symptoms reported in BBSOAS patients, in silico prediction, and in cellula effects. The mutations analyzed for the in cellula effects in this study are listed. The symptoms are categorized and simplified from the more extensive Table S1. ID = intellectual disability, DD = developmental delay, EOE/S = epilepsy and seizure, ASD = autism spectrum disorder or autism-like features, MD = motor delay, OA = optic atrophy, ODA = optic disc anomaly, NA = optic nerve anomaly, VI = visual impairment (general and CVI). * Similar predicted effects were obtained for all dimer pairs analyzed.

Based on this study, we propose the possibility of at least four different categories of BBSOAS mutations: (i) mutations associated with ID, DD, and visual pathway symptoms (e.g. VI, OA and OD/ON anomalies), such as L252P and L372P; (ii) mutations associated with ID, DD and epilepsy and ASD symptoms, such as G368D, and G395A; (iii) mutations associated with wide range of symptoms such as E318D and E400*; and (iv) mutations associated with ID, DD without visual pathway, epilepsy or ASD symptoms such as the Q244* truncation. The fact that the Q244* mutation, which was excluded from the nucleus and exerted the strongest perturbation on cellular functions tested does not cause the additional visual pathway, epilepsy, or ASD symptoms suggests the possibility that ID, DD, and MD may be associated with the loss of NR2F1 functions due to the exclusion from the nucleus or the perturbation of the function/transport of other protein partners in the cytoplasm due to dominant negative effect. Other additional symptoms may also be attributed to the modifications in the nuclear functions or NR2F1 variants, such as the changes in protein-protein interactions (dimerization partners, coactivators, corepressors), which might also act as dominant negative perturbators. The fact that the E400* mutation, which causes the loss of AF2 and thus the allosteric control of coactivator/corepressor binding and the partial exclusion from the nucleus, is associated with not only ID, DD, and MD, but also other symptoms supports this view. As an additional note, the conservative E318D mutation, which is not located in any previously described domain or interaction site but was predicted to destabilize the monomeric NR2F1, reduced NR2F1 expression and caused NR2F1, NR2F1/2 stable dimers/oligomers, is associated with a wide range of symptoms. Such observations indicate the importance of this site, which warrants more investigation. Further studies based on disease models and more clinical data will be needed to confirm our initial deductions and provide a clearer genotypic-phenotypic correlation of BBSOAS.

## DISCUSSION

The physiological function of NR2F1 as a transcriptional activator or repressor is achieved through complex protein-protein interactions involving coactivators, corepressors, and other transcription factors (Bertacchi *et al*, 2019; Montemayor *et al*, 2010; Pereira *et al*, 2000). In this study, we described the cellular and molecular consequences of BBSOAS pathogenic mutations on the function of NR2F1 with respect to its structure in solution and living cells by integrating the observations on the cellular functions with the complementary structural analysis and GCE-assisted photocrosslinking in living cells.

### Dimerization and pathogenic mutations in NR2F1

It is well described that each nuclear receptor dimer, depending on the partner and the cellular context, can trigger different regulatory events by binding to distinct sequences of target genes and controlling various biological processes (Amoutzias *et al*, 2008). Studies based on purified NR2F1 protein to examine its ability to bind DNA targets have shown that NR2F1 can bind to a variety of direct-repeat elements (DR) as a dimer (Cooney *et al*., 1992; Pinaire *et al*., 2000; Sagami *et al*, 1986; Zhang & Dufau, 2001). Such findings of purified NR2F1 set the widely referenced notion that NR2F1 exists in solution as a stable homodimer and that dimerization is a prerequisite for the DNA binding ability and function of the protein (Cooney et al., 1992). Current understanding of the role of LBD in the dimerization of nuclear receptors is based mainly on crystal structures of LBD dimers of various nuclear receptors including that of the NR2F1 homolog, NR2F2 (Kruse et al., 2008). Based on the LBD structural homology, our thorough *in silico* modeling has identified distinct mutations of amino acid residues expected to impact the LBD dimerization, some of which have then been validated in a human cell system. The *in silico* data predicted that the NR2F1 homodimer as well as the NR2F1/2 and NR2F1-RXRα heterodimers in solution would have similar contact surfaces, and that the residue G368 was critical for the homo- and heterodimerization since the G368D mutation was predicted to strongly reduce the stability and affinity of all dimeric species analyzed (Tables S2-3). This was confirmed by *in cellula* experiments.

To gain an insight into the relationship between different mutations and variable symptoms of BBSOAS patients, we compared the effects of pathogenic mutations with the WT NR2F1 in a more functionally relevant manner in cells. The clear cellular impact of the truncation mutation Q244*, where a significant portion of the LBD was lost, confirms that the LBD is required for the proper localization and cellular functions of NR2F1 in cell cycle, proliferation, and apoptosis. Interestingly, the G368D mutant was predicted to greatly reduce stability and affinity of homo- and heterodimerization while stabilizing the structure of both the auto-repressed and active monomeric forms of NR2F1, however, it had no effect on the cellular functions tested. On the other hand, the L372P mutation, predicted to strongly destabilize the monomeric structure, significantly perturbed the cell cycle, reduced cell proliferation, and increased apoptosis in a similar manner as the loss of most of the LBD by Q244* truncation (Fig. 3). Furthermore, the loss of AF2 due to the E400* truncation significantly hampered the cell cycle progression in the same manner as the Q244* truncation. Thus, our data suggest that, in a more complex cellular context, the stability of the NR2F1 monomeric structure and the interactions with other protein partners such as transcription coactivators/repressors *via* the AF2 is a more prominent factor compared to the stability and affinity of NR2F1 dimers, at least in the absence of DNA.

The question of whether NR2F1 must be dimeric in a physiological context is *per se* debated. From the early characterization of NR2F1 (Sagami *et al*., 1986), data from size-exclusion chromatography together with the ability to bind the chicken ovalbumin promoter sequence suggested that this transcriptional regulator (Wang *et al*, 1989) existed in solution as a dimer. It must be noted that the study only focused on the fraction of NR2F1 that bound the ovalbumin promoter, while other populations of NR2F1 protein were not examined. Later studies from the same group based on EMSA (Cooney *et al*., 1992) suggested that NR2F1 may bind to their response elements as a homodimer and that the ability of NR2F1 to bind a wide spectrum of response elements, including AGGTCA DRs and palindromes with various spacing was presumably due to the structural adaptation of the stable dimer.

However, some data also suggest that NR2F1 may bind to DNA as a monomer. Indeed, *in vitro* experiments suggested that NR2F1 could also bind DNA as a monomer and then dimerize (Cooney et al., 1992), albeit less efficiently. It is however worth noting that all the deductions were based on the DNA-bound NR2F1 and direct evidence showing that a stable dimer was formed prior to the addition of the DNA probes was not available. It is also possible that the pre-formed NR2F1 dimer binds preferentially to DR with short spacing, whereas in the case of regulatory elements with a variety of spacing and incomplete repeats or palindromes in a complex cellular environment, NR2F1 may bind such DNA transiently as monomer and a more stable dimer can form following the DNA-NR2F1 monomer binding (monomer pathway). Interestingly, data mining, manual curation, and data integration study reported 15 years after the early NR2F1 stable homodimer reports did not find further reports of direct evidence of functional NR2F1 homodimer (Amoutzias *et al*, 2007). Moreover, physical evidence in cellular studies that supports the notion that the stable pre-formed NR2F1 homodimer is required for its function in living cells has not been clearly demonstrated to date.

Our present data from the exogenous expression of different NR2F1 variants in cells show that the NR2F1 WT dimers could not be readily detected above the background level by NR2F1 co-IP. However, pathogenic mutations that were predicted to strongly destabilize monomeric NR2F1 (L252P, E318D, and L372P) significantly enhanced the NR2F1-NR2F1 and NR2F1/2 dimerization and/or oligomerization in cells suggesting that the pre-formed stable NR2F1 dimer/oligomer may not be a preferable state in the cellular context (Fig. 5 A-D). We propose that the formation of NR2F1 as a dimer is a very dynamic, flexible, and transient process. The presence of a stable NR2F1 homodimer, unbound to the DNA, is probably rare and may present several disadvantages for the cells as it would prevent the formation of other functional complexes between NR2F1 and other partners. It is reasonable to consider that NR2F1 may instead bind DNA individually and then recruit the second binding partner. Our data, pointing toward this notion, is supported by studies showing that even a protein with a thermodynamically stable dimerization interface does not need to form a dimer before binding DNA (Kohler & Schepartz, 2001) and in agreement with the common observation that dimeric interactions in signal transduction pathways are generally not very stable. Rather, they are dynamic and act as reversible switches in the process of the information flow (Amoutzias *et al*., 2008; Nooren & Thornton, 2003).

Notably, our data showing that the predicted impact of the mutations on the monomeric NR2F1 better correlates to NR2F1 cellular effects than the predicted impact on the DNA-free dimers also points to the likelihood that the stable pre-formed dimerization may be unnecessary for the activity of the NR2F1. This notion is also supported by studies demonstrating that the sequential monomer-binding pathway allows the protein to search for and locate a specific DNA site more quickly, resulting in greater specificity prior to equilibrium, and thus allowing a single transcription factor to recognize a number of different target sites and fine-tune the activation/repression depending on the dimerizing partners in the cells (Kohler *et al*, 1999; Kohler & Schepartz, 2001). This also aligns well with the ability of NR2F1 to adaptably bind to a number of targets and to heterodimerize with other family members, such as NR2F2 and RXR, as demonstrated here at the direct protein-protein interaction level in living cells and as previously deduced through DNA-protein complex information (Cooney *et al*., 1992; Pinaire *et al*., 2000; Rada-Iglesias *et al*, 2012; Zhang & Dufau, 2001). Indeed, the dimer formation after DNA binding has also been found for other nuclear receptor family members such as RXR, where the key molecular event leading to the cooperative enhancement of dimer formation and DNA binding is the DNA-induced conformational change that creates a favorable interface for protein-protein interactions (Holmbeck et al., 1998).

### The strength of the Genetic Code Expansion approach

Our data obtained from classical co-IP show that depending on the binding partners, different pathogenic mutations can have significantly variable effects on the capacity of NR2F1 to form stable dimers/oligomers (Fig. 5A-F). We can envisage that pathogenic mutations that favor a higher-order oligomerization of NR2F1 could interfere with the binding of NR2F1 to DNA and other coactivators/repressors and thus inhibit its proper activities. The variation of NR2F1 dimeric structures offers the diversity of target site recognition and function. Therefore, precise information on the impact of pathogenic mutations on dimerization at the cellular level is of great importance. While classical co-IP could provide some information regarding dimer formation, we could not rule out the possibility that the complex detected included not only dimers but also some higher-order oligomers. Furthermore, the lack of clear detection of WT NR2F1 dimer in the cell hampered the precise evaluation of the effect of pathogenic mutations on dimerization.

The use of site-specific GCE-enabled photocrosslinking of full-length NR2F1 in living cells has allowed us to overcome these complications and delineate the differences and similarities among the dimeric species and the pathogenic effects. Thanks to this novel approach, we were able to confirm the *in silico* prediction that the G368 residue is indeed situated in the NR2F1 homodimer and NR2F1/2 heterodimer contact sites (Fig**. 6 D-E**). Additionally, we revealed the proximity of the Q244 residue to the homodimerization and NR2F1/2 dimerization surfaces. While the similarity between the homodimer and the NR2F1/2 heterodimer is not surprising given the high LBD sequence identity, some differences including the distinct proximity of the E318 residue to the NR2F2 contact site in the heterodimer also suggest the contribution of the other domains such as the disordered N-terminal domain, which carried the most significant dissimilarity between the homologs, in the different tertiary conformations of the two dimeric species in living cells. Furthermore, our data also clearly showed that, unlike in the homodimer or NR2F1/2 heterodimer, the L372 was the most important contact residue in the NR2F1-RXRα heterodimer whereas residues Q244 and G368 were not (Fig. 6F). Based on this result, it is quite likely that the drastic L372P mutation would disrupt the NR2F1/RXRα dimer as predicted by our structural analysis. Meanwhile, we also observed that the Q244 residue was also in the proximity of the NR2F1-RXRα dimerization site (Fig. 7C). However, it was clearly much less involved than the L372 residue (Fig. 6F).

Our further analysis of the effects of pathogenic point mutations using the crosslinking AzF at Q244 position showed that the G368D mutation resulted in a significant decrease in both homo- and heterodimerization (**Fig. 7 A-C**), confirming the *in silico* prediction that this mutation caused the drastic destabilization and the loss of affinity of all the dimeric pairs analyzed. Interestingly, the distinct reduction of NR2F1/2 dimerization observed for the G395A mutation and not paralleled by consistent *in silico* predictions regardless of the auto-repressed or active state of NR2F1 LBD not only demonstrates the divergence among the conformation of different dimer pairs but also suggests the contribution of AF2 in the NR2F1/2 heterodimerization. Also of note is that the important mutation at the NR2F1-RXRα contact site, L372P, which was expected to significantly disrupt NR2F1/RXRα dimerization, did not impact the capture of the dimer via Q244AzF. This was not surprising since the difference in the tertiary structure between the NR2F1/RXRα and the NR2F1 homodimer or NR2F1/2 dimer in living cells was clear (Fig. 6D-F). Some changes in the dimer conformation NR2F1/RXRα may not have been captured as efficiently via Q244AzF.

Overall, these data highlight the diversity of quaternary conformations of full-length NR2F1 in contact with different protein partners in living cells and complement the *in silico* structural prediction derived from the protein in solution. Noteworthy is that the predicted changes in affinity and stability of the NR2F1 dimer attributed to the pathogenic substitutions are highly correlated (R^2^ = 0.807 for NR2F1 homodimer in heterozygosis). The significant correlation between mutation-induced variations in dimerization affinity and folding stability in the LBD suggests that the pathogenic mechanisms underlying BBSOAS may also involve a synergy between the LBD dimerization process (pre-/post-DNA binding) and its functional properties.

### Discovery of CRABP2-NR2F1 binding *via* the coactivator recognition site

Our data suggest that the dysfunction of NR2F1 in the cellular context correlates well with the disruption of the monomeric structure integrity due to the L252P mutation in the CRS and the loss of AF2 and thus the allosteric control on the protein interaction at the CRS due to the E400* truncation. To date, information regarding the interacting partners of NR2F1 at the CRS is lacking. The fact that in living human cells NR2F1 heterodimerized with RXR, the key RA-induced transcription regulator (Evans & Mangelsdorf, 2014), and that NR2Fs either antagonize retinoid-dependent gene expression (Neuman *et al*, 1995), or are themselves modulated by retinoids (Alfano *et al*, 2014; Clotman *et al*, 1998; Tran *et al*, 1992; Zhuang & Gudas, 2008), suggest that NR2F1 might be a key regulator of the retinoid signaling pathway during embryonic development by playing distinct functional roles in a context- and time-dependent manner (Alfano *et al*., 2013). Moreover, overexpression of some of the NR2F1 variants, in particular, the Q244* and the L252P mutations, which lead to abnormal localization in the cytoplasm (Fig. 4), led us to explore the interaction between NR2F1 and CRABP2, a plasmonuclear shuttling protein, which transports RA to the nucleus and is known to act as a coactivator of RAR (Sessler and Noy, 2005).

The CRABP2-RAR complex is a short-lived intermediate (Budhu & Noy, 2002; Dong *et al*, 1999). We expected that if the CRABP2-NR2F1 interaction existed, it would have eluded the detection by conventional methods used for studying NR2F1 protein-protein interaction so far. And this was indeed the case. Thanks to the photocrosslinker placed at the CRS (Q244AzF), we could demonstrate for the first time a direct binding of CRABP2 to the CRS of NR2F1 (Fig. 8A). Our structural analysis (Fig. 8E) also supports our findings and the notion that, in cells, CRABP2 may channel RA to NR2F1 in a similar manner as the RA transfer to RAR (Budhu and Noy, 2002). Furthermore, we observed that different pathogenic mutations exerted distinct effects on the ability of NR2F1/CRABP2 complex formation. The expression of mutations that strongly destabilize the monomeric structure of NR2F1, namely, the L252P in the CRS and L372P at the RXR-NR2F1 contact site abrogated the NR2F1/CRABP2 complex, again highlighting the monomeric structure integrity of NR2F1 as an important predicting factor of NR2F1 protein-protein interaction. It should also be noted that based on structural alignment with RAR (3KMR) the L252 residue of NR2F1 along with the CRS overlap the predicted binding function-3 (BF-3) of RAR which is highly conserved among nuclear receptors but has not been extensively characterized(Buzón et al., 2012).

The binding of CRABP2 to CRS/BF-3 of NR2F1 further supports the notion that the functional relationship between the two proteins may be similar to that between CRABP2 and RARL. On the other hand, the inverse results showing that the CRABP2-NR2F1 binding was strongly enhanced by the G368D mutation, which reduced NR2F1 homodimerization, NR2F1/2 and NR2F1-RXRα heterodimerization and the G395A mutation which reduced NR2F1/2 homodimerization, suggest a scenario where the loss of dimerization, especially of NR2F1 homodimerization and NR2F1/2 dimerization may tip the balance toward CRABP2-NR2F1 binding and possibly the RA-induced transcription pathway. Notably, the G395 residue locates between the CRS/BF-3 and the AF2 modules. The aberrantly high CRABP2-NR2F1 interaction due to the G395A mutation may be attributed to the loss of allosteric control of the CRS/BF-3 by AF2. The fact that most of the pathogenic mutations studied here significantly interfered with the CRABP2 interaction with NR2F1 infers the physiological importance of this interaction. It has been shown that the transcription of NR2F1 and CRABP2 are both induced by RA in mouse embryonic stem cells (Quintero *et al*, 2018), suggesting context-dependent co-expression of the two proteins. Indeed, our immunofluorescence of the anterior part of the forebrain and the eye region of mouse embryos showed that NR2F1 and CRABP2 selectively co-expressed in the forming hippocampus and ventral retina, supporting the physiological importance of the CRABP2-mediated NR2F1 function in the brain, particularly in the context of hippocampal and visual pathways.

Together, our data demonstrate the allosteric relationship between the less known but highly conserved surface on the LBD of nuclear receptors, CRS/BF-3, and the AF2 and the consequence of pathogenic mutations therein on the interaction between NR2F1 and its newly identified partner, CRABP2, and propose the possibility that RA may function as NR2F1 ligand in some physiological contexts.

## CONCLUSION

A crucial piece of information to help clinicians design proper treatment plans and predict disease progression is to establish a clear genotype-phenotype correlation among patients. This requires knowledge of the structure-function relationship of the pathogenic protein.

By combining computational analysis of protein structure and the power of genetic code expansion technology to greatly improve the sensitivity and specificity of protein-protein interaction study, we present new insight into the structure-function relationship of NR2F1 in the context of BBSOAS pathology. The structural analysis of the isolated and dimerized LBD provides an understanding of the molecular interaction between the LBD of NR2F1 and its partners without the complex interactions from other cellular partners, whereas the complementary GCE-enabled site-specific photocrosslinking in living cells further highlights the variable quaternary conformations of NR2F1 with functional relevance in a more complex setting of the cell, where the contribution of different domains of the full-length protein and interaction with other cellular actors were also taken into account. Our data shed light on the scenarios that may connect the variable BBSOAS symptoms to different point mutations in the NR2F1 LBD.

The complementary approaches described here not only provide information that will guide further studies to arrive at a better understanding of the genotype-phenotype relationship presented in BBSOAS but also can be applied to a wide range of studies involving protein structure-function relationships presented in other genetic diseases.

## ACKNOWLEDGMENTS and FUNDING

We thank the Genetic Code Expansion facility at iBV for the assistance in applying the GCE technology. We also thank A. Loubat at the flow cytometry facility at iBV. This work was funded by the French Government (National Research Agency, ANR) through the ‘Investments for the Future’ programs IDEX UCAJedi ANR-15-IDEX-01, by the “Fondation de la Recherche Médicale (Equipe FRM2020)” (#EQU202003010222), “Fondation de France” (# 00123416/WB-2021-38751), ERA-NET Neuron grant (Brain4Sight) (ANR-21-NEU2-0003-03) grants to M.S. V.M was the recipient of a research contract within the FSE REACT EU-PON R&I 2014-2020 granted to D.D.O.).

## MATERIALS and METHODS

### Molecular modeling of NR2F1 Ligand-binding domain (LBD) in its auto-repressed and active conformations

The three-dimensional structure of human NR2F1 LBD in its auto-repressed conformation was obtained by homology modeling using the crystal structure of human NR2F2 LBD as a template (PDB entry 3CJW, resolution 1.48 Å (Kruse et al., 2008), which shares 96% sequence identity with that of NR2F1. The missing loops encompassing residues 201-213 and 276-292 were modeled using Maestro (Schroedinger). The structure of human NR2F1 LBD in its active conformation was modeled as previously elucidated in (Khalil et al., 2022), thus using the structure of RXRα in complex with 9-cis-retinoic acid (9cRA) as a template (PDB entry 1FM6 (Gampe et al., 2000), ∼40 % sequence identity with human NR2F1 LBD) after removal of the co-activator peptides. The resulting model of active NR2F1 LBD was then employed for docking and mutagenesis analyses. All molecular models were prepared following the *Protein Preparation Wizard* pipeline included in Bioluminate (Schroedinger), which provides for assigning bond orders according to the Chemical Components Dictionary database (www.pdb.org, wwPDB Foundation, Piscataway, NJ, USA), adding H atoms, selecting the most probable rotamer in case of alternative conformations of the sidechains and modeling of the missing loops. The structures were then refined by sampling the orientation of water molecules and predicting the protonation states of ionizable amino acids at pH 7.5 using PROPKA (Li *et al*, 2005) to assign and optimize H-bonds. The last step of protein preparation consisted of the minimization of the model structures using OPLS4 forcefield (Schroedinger, New York, NY, USA) using 0.3 Å as threshold for the Root-Mean Square Displacement (RMSD) of heavy atoms.

For *in silico* docking simulations, the three-dimensional structural model of human NR2F2 LBD was obtained by applying the *Protein Preparation Wizard* pipeline to the PDB entry 3CJW (Kruse et al., 2008), where missing residues 194-206 and 269-285 were modeled by Maestro (Schroedinger). Human RXRα was modelled by following the *Protein Preparation Wizard* pipeline for the respective molecule present in PDB entry 4NQA.

### Docking simulations of NR2F1 LBD in its auto-repressed and active state

To establish a putative NR2F1 interaction model in native-like conditions, we superimposed the experimental structures of the dimeric assemblies of homologous nuclear receptors, namely NR2F2 homodimer (PDB entry 3CJW (Kruse et al., 2008)), RXRα homodimer (PDB entry 1MZN (Egea et al., 2002)), RXRα-Retinoic Acid Receptor β (RARβ) heterodimer (PDB entry 5UAN (Chandra et al., 2017) and RXRα-LXRβ heterodimer (PDB entry 4NQA (Lou et al., 2014)). All experimental dimers presented highly similar and conserved (Figure S2) interfaces in terms of the physicochemical properties of interacting residues (Figure S3), therefore the dimeric assembly corresponding to NR2F2 homodimer was selected as the native-like conformation. Rigid-body docking simulations of NR2F1 LBD with its putative partners, namely NR2F1 (homodimer), NR2F2 and RXRα LBDs, were performed using PIPER [16933295] module implemented in Bioluminate (Schroedinger), by setting 70000 as the number of tested poses with a 5° sampling of Euler angles. Modeled loops 201-213, 276-292 and the C-terminal loop (residues 407-414) of both auto-repressed and active NR2F1 LBD were excluded from the docking interface to avoid potential artifacts due to highly flexible regions. For the same reason, modeled loops 193-207, 268-286, and 400-407 of human NR2F2 LBD were masked during docking simulations. The best 1000 resulting poses from docking simulations were then grouped in ∼100 clusters, whose centroids were aligned against the native-like conformation to identify the native-like cluster with the lowest RMSD with respect to PDB entry: 3CJW (NR2F2 homodimer). Finally, the highest-scored conformation of the native-like cluster was selected as the model for WT NR2F1 LBD-partner interaction. Each of the eight obtained complexes (NR2F1 homodimer in homo and heterozygosis, NR2F1-NR2F2 heterodimer and NR2F1-RXRα heterodimer, with NR2F1 in both its auto-repressed and active form) underwent the *Protein Preparation Wizard* pipeline prior to *in silico* mutagenesis with the same parameters as detailed in the previous section.

The modeled NR2F1 LBD was also subjected to rigid-body docking using ZDOCK 2.3 (Chen et al., 2003) to evaluate homodimer formation propensity based on the correlation between ZDOCK score and experimentally determined binding constants (Dell’Orco et al., 2007). We carried out 4 rigid-body docking simulations for both the auto-repressed and the active form of NR2F1 LBD, starting from 4 different relative orientations of the monomers with dense sampling (6° sampling step), yielding 4000 potential docking poses for each simulation. The final 16000 poses were filtered to identify those resembling the NR2F1 dimer obtained with PIPER, by setting the threshold of the Cα-RMSD to 1 Å.

### Molecular modeling of CRABP2 and docking simulations with active NR2F1 LBD

The structures of apo and holo human CRABP2 (unbound/bound to retinoic acid) were retrieved from PDB entries 2FS6 and 2FS3, respectively (Vaezeslami et al., 2006). Before docking simulations, both CRABP2 models underwent the same *Protein Preparation Wizard* procedure as elucidated in the previous sections. Unbiased docking simulations of active NR2F1 LBD with CRABP2 were carried out using PIPER (Kozakov et al., 2006) module implemented in Bioluminate (Schroedinger), by setting 70000 as the number of tested poses with a 5° sampling of Euler angles.

The best 1000 resulting poses from docking simulations were then grouped in ∼30 clusters, whose centroids were filtered according to the simultaneous presence of CRABP2 residues Q75, P81, and K102 (whose mutations compromise binding of CRABP2 to RXRα (Budhu and Noy, 2002) and NR2F1 residue Q244 (used for GCE crosslinking) in the NR2F1-CRABP2 interface. Finally, the Cα of the 4 filtered solutions (2 for apo and 2 for holo CRABP2) obtained from docking simulations were superimposed on each other to identify the potential native-like conformation (RMSD = 2.123 Å) in which apo and holo CRABP2 essentially shared the interaction interface with NR2F1 regardless of the presence of the retinoid.

### *In silico* prediction of relative stabilities and affinities of NR2F1 LBD missense variants

All missense mutations in NR2F1 LBD monomer, homo- and heterodimers were introduced using the *Residue Scanning* tool from Bioluminate (Schroedinger); for NR2F1 homodimer mutations were generated in both homo- and heterozygosis. The most probable rotamer for the sidechain of each aminoacidic substitution was automatically assigned before performing the same energy minimization protocol as above. We estimated the effects of each substitution on the Gibbs free energy of folding (ΔΔG_f_^app^ = ΔG_f_^app,mut^ - ΔG_f_^app,WT^) of both the auto-repressed and the active forms of isolated NR2F1 LBD and its complex with NR2F1 (both in homo- and heterozygosis), NR2F2 and RXRα compared to the wildtype. The computation of ΔΔG_f_^app^, expressed in kcal/mol, was performed by calculating the variant-specific thermodynamic cycle using the Molecular Mechanics/Generalized Born and Surface Area Continuum solvation (MM/GBSA) method, which does not include the explicit energetic contribution deriving from conformational changes. Therefore, the ΔΔG_f_ ^app^ reported in Supplementary Tables 1-3, should not be considered as precise thermodynamic quantities, but rather “apparent” values. Positive values of ΔΔG_f_ ^app^ indicate a destabilizing effect for the specific mutation, whereas negative values identify a stabilizing mutation. The variations in Gibbs free energy of binding (ΔΔG_b_^app^ = ΔG_b_^app,mut^ - ΔG_b_^app,WT^) reported in Supplementary Tables 2 and 3 were calculated between NR2F1-LBD and the partner nuclear receptor (NR2F1 in homo/heterozygosis, NR2F2 and RXRα). Positive values of ΔΔG_b_^app^ indicate decreased affinity for the specific ligand, whereas negative values identify stronger binding.

### Molecular Dynamics simulations of NR2F1-LBD nonsense mutations

Molecular Dynamics simulations of isolated human NR2F1-LBD variants were set up based on the model of the auto-repressed form described in the previous sections. The models for nonsense variants E400* and Q244* were generated by truncating the structure of the WT after the carbonyl group at the C-term of the respective residue and capping both N- and C-terms with a NH_2_ group. All protein systems underwent the same *Protein Preparation Wizard* procedure described in previous sections before setting up the simulation system. All-atom MD simulations of NR2F1-LBD variants were run on GROMACS 2016.5 (Abrahams *et al*, 2013) simulation package, setting CHARMM36m (Huang *et al*, 2017) as forcefield. The size of the simulated protein systems was ∼45000 atoms for WT and E400* variants and ∼30000 atoms for Q244* mutant, systems were prepared and minimized according to the protocol and parameters detailed in (Marino *et al*, 2015). Briefly, proteins were put in the center of a dodecahedral box with the edges set 12 Å apart from any protein atom to avoid potential interactions with the periodic images, then the systems were neutralized with 150 mM KCl and underwent a two-step energy minimization procedure with steepest descent and conjugate gradients algorithms. System equilibration was carried out for 2 ns in NVT ensemble (T = 310 K) with position restraints followed by 2 ns with no position restraints as explained in (Marino & Dell’Orco, 2016), while productive 500 ns runs were carried out in NPT ensemble (T = 310 K, P = 1 atm) using the setup elucidated in (Marino & Dell’Orco, 2019). The Cα-RMSD with respect to the protein after equilibration was calculated using *gmx rms*, while the time-averaged Cα-RMSD over the 500 ns trajectory with respect to the average structure, that is the Root-Mean Square Fluctuation (RMSF), was calculated by *gmx rmsf*. To evaluate the conformational rearrangement due to the chain truncations, we calculated the angles between helices 1 (identified by the vector connecting the Cα of C183 and R194) and 2 (identified by the vector connecting Cα of C220 and R235) and between helices 1 and 8 (identified by the vector connecting Cα of S382 and F390) by means of *gmx gangle*. All *gmx* functions were provided by GROMACS 2016.5 simulation package.

### Cell preparation and transfection

The HEK293T cell line was maintained at 37°C in a humidified incubator with 5% CO_2_ in DMEM supplemented with 10% fetal bovine serum (FBS). The cells were seeded at 350,000 cells per well in 6-well plates for oligomerization and dimerization experiments, 150,000 cells per well in 12-well plates to study the cell cycle and apoptosis, and 80,000 cells per well in 24-well plates for immunofluorescent staining. After 24 hours, the cells were transfected with JetPRIME transfection reagent with the 1.5, 1, 0.1875 ug total DNA amount for 6-, 12-, and 24-well plates respectively.

### Plasmids

The plasmid for the expression of the C-terminally FLAG-tagged human NR2F1, pcDNA3.1-NR2F1-DYK (Genscript clone ID: OHu23866D), was used as the template for the site-directed mutagenesis (Liu & Naismith, 2008) to produce NR2F1 variants with amber codon and pathogenic mutations. To generate the constructs to express Myc-tagged protein partners, i.e., pcDNA-Myc-NR2F1, pcDNA-Myc-NR2F2, and pcDNA-Myc-RXRα, EcoRI/XhoI restriction sites were added to flank the *NR2F1*, *NR2F2*, and *RXRα* genes in pcDNA3.1-NR2F1-FLAG, pME-NR2F2, and pcDNA3.1-hRARα.hRXRα, respecitively. The genes were then excised and cloned to replace DBC1 in the pcDNA Myc DBC1 plasmid by restriction ligation at the EcoRI/XhoI restriction sites. pME-NR2F2 was a gift from Nathan Lawson (Addgene plasmid #138359; http://n2t.net/addgene:138359; RRID:Addgene_138359). pcDNA Myc DBC1 was a gift from Osamu Hiraike (Addgene plasmid # 35096 http://n2t.net/addgene:35096; RRID:Addgene_35096). pcDNA3.1-hRARα.hRXRα was a gift from Catharine Ross (Addgene plasmid #135411; http://n2t.net/addgene:135411; RRID:Addgene_135411). pIRE4-Azi was a kind gift from Irene Coin (Addgene plasmid #105829; http://n2t.net/addgene:105829; RRID:Addgene_105829).

### Incorporation of AzF and photo-crosslinking

Twenty-four hours before the transfection, cells were seeded at 350,000 cells per well in 6-well plates. Using the JetPRIME transfection reagent (Polyplus), cells were co-transfected with pIRE4-AziRS and pcDNA3.1-NR2F1-FLAG (WT or variants) plasmids at the ratio of 3:1, respectively. For dimerization experiments, the plasmids carrying Myc-NR2F1, NR2F2, or RXRα were transfected at the same amount as FLAG-NR2F1. A fresh 150-mM p-azido-l-phenylalanine (AzF) solution was prepared on the day of the transfection by dissolving the AzF powder (Santa Cruz) in 0.375N NaOH and 25% DMSO. A 3-mM working solution (WS) of AzF in a complete medium supplemented with 100 mM HEPES was prepared and added to the well to obtain the final AzF concentration of 300 μM. Forty-eight hours after the transfection, the cells washed twice with DMEM to remove AzF and the medium was replaced with the DMEM containing 10% FBS and 10 mM HEPES. Protein photocrosslinking was performed by irradiating the cells with 365-nm UV for 15 minutes under a UV lamp (UVITEC LF-215.LS LAMP 365/254NM 2X15W 230V).

### Immunoprecipitation

Cells were washed with PBS and lysed with prechilled RIPA buffer (50 mM Tris-HCL, pH 8.0 with 150 mM sodium chloride, 1.0% NP-40, 0.5% sodium deoxycholate and 0.1% sodium dodecyl sulfate) with cocktail protease inhibitor (cOmplete^TM^, Sigma-aldrich) for 30 minutes in at 4°C on a shaker. Lysed cells were scraped off the plate and transferred to 1.5 μl Eppendorf tube and sonicated for 5 seconds on ice. Cells debris were removed by centrifugation (14,000*x*g, 10 minutes, 4°C). The supernatants were immunoprecipitated using ANTI-FLAG® M2 Affinity Gel (Sigma-Aldrich) or Sepharose beads coupled with the anti-CRABP2 antibody (Proteintech) overnight at 4°C. The beads were collected by centrifugation (5,000*x*g, 1 minute, 4°C) and washed three times with TBS. The immunoprecipitated proteins were eluted from the beads in LDS sample buffer (NuPAGE™ LDS Sample Buffer, NP0008, ThermoFisher) by heating for 10 minutes at 70°C. The eluted proteins were separated from the beads by centrifugation (5,000*x*g, 1 minute) and transferred to 1.5 μl Eppendorf tube. Proteins were denatured by dithiothreitol at the final concentration 0.1 M.

### Western blot

The samples were migrated in the 6-15% gradient polyacrylamide gels and blotted onto the PVDF membrane (Amersham Hybond P 0.45 PVDF blotting membrane, Cytiva). The membranes were probed using the primary antibodies, i.e., mouse anti-Myc tag (Cell Signaling Technology), rabbit anti-DDDDK (FLAG) tag (Genetex), rabbit anti-histone H3 (R&D), rabbit anti-CRABP2 (Proteintech), and peroxidase-conjugated anti-rabbit or anti-mouse secondary antibodies. The membranes were detected using a western blot detection reagent (ECL™ Prime Western Blotting System, Cytiva).

### Immunofluorescence microscopy

After 48 hours of transfection, HEK293T cells were fixed with 4% paraformaldehyde (PFA) for 15 minutes at room temperature (RT) and then washed 3 times with PBS. The cells were pre-blocked with pre-blocking solution (PBS, 0.3% tween, 5% serum) for 1 hour at RT with gentle shaking every 15 minutes. Cells were stained with primary antibodies; rabbit anti-NR2F1 (1:1000) (Abcam) and mouse anti-acetylated-tubulin (1:1000) (Sigma) for 4 hours at RT with gentle shaking every hour, followed by washing with PBS containing 1% serum. After the wash, cells were stained with The secondary antibodies, Alexa Fluor-488 anti-rabbit and Alexa Fluor-647 anti-mouse antibodies and DAPI for 1 hour at RT and washed 3 times with PBS. Cells were mounted on glass coverslips with the mounting medium (PBS, 2% N-propyl gallate, 90% Glycerol) and analyzed using Zeiss 710 confocal microscope equipped with a 405nm diode, an argon ion, a 561nm DPSS, and 647 HeNe lasers using a 40X objective with oil immersion.

### Immunostaining for cell cycle and apoptosis assays

After 48 hours of transfection with pcDNA-NR2F1-FLAG (WT or pathogenic variants), HEK293T cells were fixed with 70% cold ethanol for 15 minutes and then washed with 3 ml of PBS containing 1% FBS. The fixed cells were blocked with a pre-blocking solution (PBS, 0.3% tween, 5% serum) and then incubated for 30 minutes at 4°C with gentle shaking. The cells were co-stained with primary antibodies, mouse anti-NR2F1 (1:1000) (R&D), rabbit anti-phospho-Histone H3 (pH3) (1:1000) (Millipore), or rabbit anti-cleaved Caspase 3 (1:1000) (Cell signaling) and incubated for 2 hours at 4°C with gentle shaking. After washing, cells were stained with Alexa Flour 488 anti-mouse and Alexa Flour 647 anti-rabbit antibodies for 1 hour at 4°C with gentle shaking. Then the cells were washed with PBS. To study the cell cycle, cells were stained with propidium iodide (PBS, 0.1% NP40, 0.2% RNase, 3.95% propidium iodide) for 15 minutes at RT followed by 2 washes with PBS. After filtering through a 70-micron filter, the cells were analyzed by FACS.

### Immunofluorescence of mouse brain

All mouse experiments were conducted in accordance with the relevant national and international guidelines and regulations (European Union rules; 2010/63/UE), and with approval by the local ethical committee in France (CIEPAL NCE/2019–548). Standard housing conditions were approved by the local ethical committee in France (CIEPAL NCE/2019–548). Briefly, adult mice were kept on a 12-h light–dark cycle and three animals were housed per cage with the recommended environmental enrichment (wooden cubes, cotton pad and igloo), and with food and water *ad libidum*. Whole heads of embryonic (E) 13.5 mouse embryos were dissected on ice-cold PBS 1X and fixed in 4% paraformaldehyde (PFA) at 4°C for 3 hours in agitation, then washed in PBS 1X and dehydrated in 25% sucrose overnight at 4°C. Heads were then embedded in optimal cutting temperature compound (OCT) and stored at −80°C. Cryostat sections (12-14 μm) were collected on SuperFrost slides and subjected to immunofluorescence, as previously described (Harb *et al*, 2021). Briefly, the brain sections were washed and unmasked in sodium citrate 85 mM (pH 6)., 95-100°C for 10 minutes. After pre-blocking, the samples were stained with rabbit anti-CRABP2 (102251-AP; Proteintech) and mouse anti-NR2F1 (H8132; R&D) antibodies followed by corresponding secondary antibodies and counterstaining with DAPI. After subsequent washing and mounting on slides, the samples were analyzed using an Apotome Zeiss microscope with a 20X objective, and with the AxioVision Software.

### Statistical analysis

Statistical analyses (ANOVA or t-test) were performed using GraphPad Prism9 and data are presented as mean ± SD. N values represent biological replicates from at least 3 independent experiments.

## SUPPLEMENTARY FIGURES AND TABLES LEGENDS

**Figure S1. Multiple sequence alignment of NR2F1, NR2F2, LXRβ, RXRα and RARβ together with the consensus sequence (cons).** The disordered and hinge regions are represented by a line and labeled, the DBD is identified by the green box, the LBD is identified by the light blue box, residues whose variants are associated with BBSOAS are shown in bold and red. The region of the sequence where residues belonging to the dimerization interfaces (DI) lie in all four proteins is shaded in orange, Co-activator Recognition site (CRS) in purple, Activation Function 2 (AF2) helix in red.

**Figure S2. Dimerization interfaces of the template complexes used to infer the potential dimerization interface of NR2F1, namely NR2F2 homodimer (PDB entry: 3CJW), RXRα homodimer (PDB entry: 1MZN), RXRα-Retinoic Acid Receptor β (RARβ) heterodimer (PDB entry: 5UAN) and RXRα-LXRβ (PDB entry: 4NQA) heterodimer.** The molecular surface of NR2F2 protomers A and B is shown in gray and cyan, respectively; the molecular surface of RXRα protomers A and B is shown in light pink and light blue, respectively; the molecular surface of LXRβ is shown in yellow, the molecular surface of RARβ in light green. Residues belonging to the dimerization interface are shown as sticks with N atoms in dark blue, O atoms in red, and C atoms depending on the physicochemical nature of the interaction: electrostatic (blue), hydrophobic (green), H-bonds (orange). Bottom panels show all the residues belonging to the dimerization interfaces (in sticks) grouped by their physicochemical properties. Purple sticks represent NR2F1 interfacial residues identified by docking simulations.

**Figure S3. FACS gating for HEK293T cell cycle and apoptosis analysis.** (**A**) Cell cycle phases (G0/G1, S, G2/M) of NR2F1-WT expressing HEK293T cells were analyzed using propidium iodide staining. This figure shows the number of cells in different cell cycle phases. (B) The NR2F1-positive population (green) is distinguished from the NR2F1-negative population (blue) by NR2F1 staining. (C) Cells in the mitotic phase are indicated in the square (M) by PH3 staining along with propidium iodide staining. (D) Cleaved-caspase 3 was used to indicate apoptotic cells. NR2F1-positive apoptotic cells are cleaved-caspase3 and NR2F1 double-positive and are indicated in the upper right quadrant.

**Figure S4. Morphology of HEK293T cells expressing the truncated NR2F1-Q244* protein.** After 48 hours of transfection of NR2F1-WT (**A**) and NR2F1-Q244* (**B**), cells were observed under a bright-field microscope. The lower number of cells expressing NR2F1-Q244* was observed with a significant portion of cells exhibiting the morphology corresponding to apoptotic cells.

**Figure S5. Site-specific photo-crosslinker incorporation in the LBD of NR2F1 and covalent protein complex capturing in living HEK293T cells.** AzF incorporation was performed as described in Figure 6 and Methods. Top: Immunoblots show the AzF incorporation after 48 hours of transfection. Truncated NR2F1 (boxed) was produced in the absence of AzF. Once AzF was present in cultural media, the full-length NR2F1 (arrow) was produced by amber codon suppression. The large protein complexes (*) can be observed where NR2F1-AzF protein was produced and photo-crosslinked by UV irradiation (365 nm). Bottom: Immunoblot showing different molecular weights of truncated forms which varied depending on the positions of the amber codon placed in the LBD. The difference in the molecular weights of the photo-crosslinked complexes of NR2F1 proteins that carried AzF at different positions suggests the difference in the protein partner identity and/or in the conformation of the crosslinked complex.

**Figure S6. Full-length NR2F1-Q244AzF functions equivalently to the NR2F1-WT in HEK293T cells.** and The quantification of mitotic (**A**) and apoptotic (**B**) cells was carried out by FACS analysis. The comparison between cells expressing the truncated NR2F1-Q244* and the full-length NR2F1-Q244AzF proteins indicated that the incorporation of AzF to yield the full-length NR2F1-Q244AzF restored to the cells the ability to proliferate and rescued the cells from apoptosis to the levels of cells expressing NR2F1-WT. (**C**) Immunostaining of NR2F1 (red) shows that the truncated NR2F1-Q244* localizes in the cytoplasm (co-localization with tubulin, green). The AzF incorporation to yield the full-length NR2F1-Q244AzF protein restores the nuclear localization in the same manner as NR2F1-WT (co-localization with DAPI staining of the nucleus, blue).

Video S1. Three-dimensional structure of NR2F1-DNA complex. Protein and DNA structures are represented as a cartoon, with the LBD shown in green, the DBD in cyan, and the dsDNA in purple.

Video S2. The three-dimensional structure of the LBD is shown as a cyan cartoon with the AF2 helix shown in red and the CRS in green. The Cα of the residues whose mutations are associated with BBSOAS is represented as yellow spheres and labeled.

Video S3. Three-dimensional structure of NR2F1 LBD. Protein structure is shown as a cyan cartoon, the AF2 helix is shown in red and the CRS in green. The Cα of the residues whose mutations are associated with BBSOAS is represented as spheres, labeled, and colored in a red-yellow-green scale according to the ΔΔGfapp values of the auto-repressed NR2F1 homodimer in heterozygosis reported in table ST1.

**Table S1. Summary list of NR2F1 variants in the LBD and clinical description of BBSOAS reported patients.** BBSOAS patients, identified by their protein variant and—when available—by their LOVD identifier, are listed following the chronological order of reports and publications describing their cases. Main clinical signs include altered brain morphology as observed by MRI, developmental delay (DD), intellectual disability (ID), visual system deficits, early-onset epilepsy and seizures (EOE/S), autism spectrum disorder (ASD), and behavioral abnormalities and hypotonia. An extended version of these data can be found in Bertacchi et al., 2022. Abbreviations and reference lists are listed below the table.

**Table S2. Predicted effects of BBSOAS-associated variants of NR2F1 LBD on the stability (ΔΔGfapp) of the isolated auto-repressed NR2F1 LBD and on both the stability and affinity (ΔΔGbapp) of the auto-repressed NR2F1 LBD in complex with LBD from NR2F1, NR2F2 and RXRα.** For NR2F1 homodimers, the effects of the mutations were evaluated in both heterozygosis (het) and homozygosis (hom) genetic conditions. ΔΔG values are expressed in kcal/mol and are highlighted in a color scale red-yellow-green from the most to the least detrimental to the respective property. Negative values indicate a stabilizing effect of the mutation.

**Table S3. Predicted effects of BBSOAS-associated variants of NR2F1 LBD on the stability (ΔΔG_f_^app^) of the isolated active NR2F1 LBD and on both the stability and affinity (ΔΔG_b_ ^app^) of the active NR2F1 LBD in complex with LBD from NR2F1, NR2F2 and RXRα.** For NR2F1 homodimers, the effects of the mutations were evaluated in both heterozygosis (het) and homozygosis (hom) genetic conditions. ΔΔG values are expressed in kcal/mol and are highlighted in a color scale red-yellow-green from the most to the least detrimental to the respective property. Negative values indicate a stabilizing effect of the mutation.

**Table S4. Results from rigid-body docking performed with ZDOCK 2.3.** ^a^Number of docked complexes with a Cα-RMSD < 1 Å with respect to the native structure obtained by docking simulations performed with PIPER. ^b^Rank of the native-like docked pose with the highest score among the 4000 poses of each of the four docking replicas. ^c^ZDOCK-score of the native-like solutions reported as average ± standard deviation.

**Table S5. List of primers used in this study.**

**Table S6. List of antibodies used in this study.**

