## Supplementary Information for "Integrated genetic code expansion and structural bioinformatics reveal disrupted supramolecular assembly in a genetic disorder"

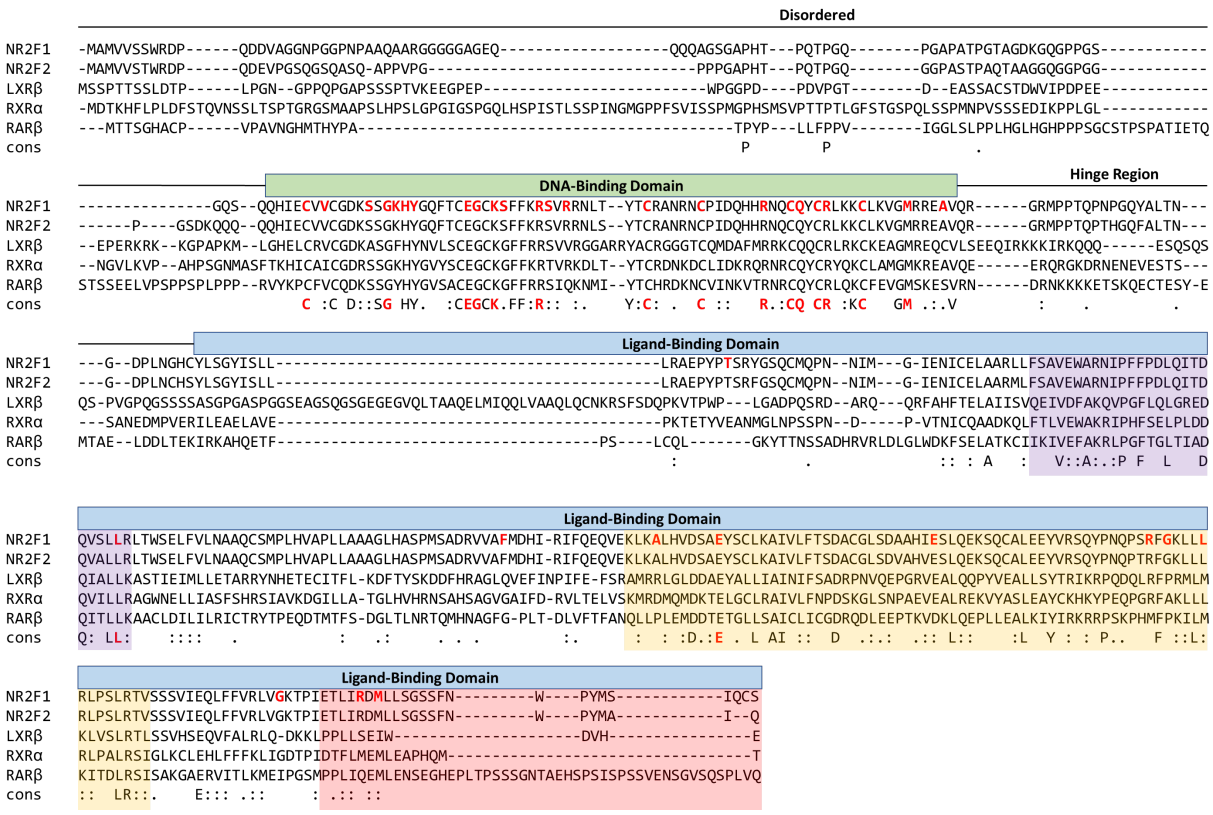


***Figure S1. Multiple sequence alignment of NR2F1, NR2F2, LXRβ, RXRα and RARβ together with the consensus sequence (cons).*** *The disordered and hinge regions are represented by a line and labeled, the DBD is identified by the green box, the LBD is identified by the light blue box, residues whose variants are associated with BBSOAS are shown in bold and red. The region of the sequence where residues belonging to the dimerization interfaces (DI) lie in all four proteins is shaded in orange, Co-activator Recognition site (CRS) in purple, Activation Function 2 (AF2) helix in red.*

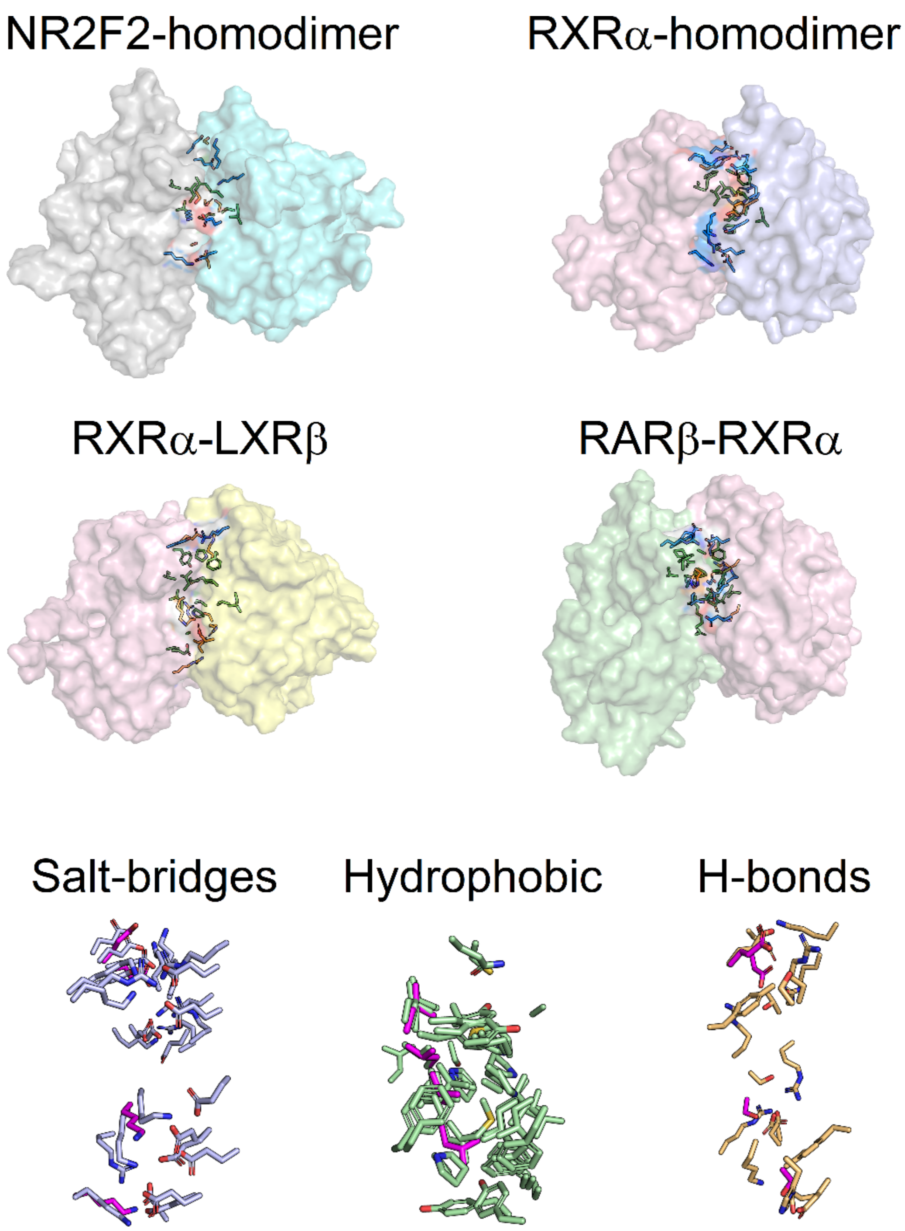


***Figure S2. Dimerization interfaces of the template complexes used to infer the potential dimerization interface of NR2F1, namely NR2F2 homodimer (PDB entry: 3CJW), RXRα homodimer (PDB entry: 1MZN), RXRα-Retinoic Acid Receptor β (RARβ) heterodimer (PDB entry: 5UAN) and RXRα-LXRβ (PDB entry: 4NQA) heterodimer.*** *The molecular surface of NR2F2 protomers A and B is shown in gray and cyan, respectively; the molecular surface of RXRα protomers A and B is shown in light pink and light blue, respectively; the molecular surface of LXRβ is shown in yellow, the molecular surface of RARβ in light green. Residues belonging to the dimerization interface are shown as sticks with N atoms in dark blue, O atoms in red, and C atoms depending on the physicochemical nature of the interaction: electrostatic (blue), hydrophobic (green), H-bonds (orange). Bottom panels show all the residues belonging to the dimerization interfaces (in sticks) grouped by their physicochemical properties. Purple sticks represent NR2F1 interfacial residues identified by docking simulations.*

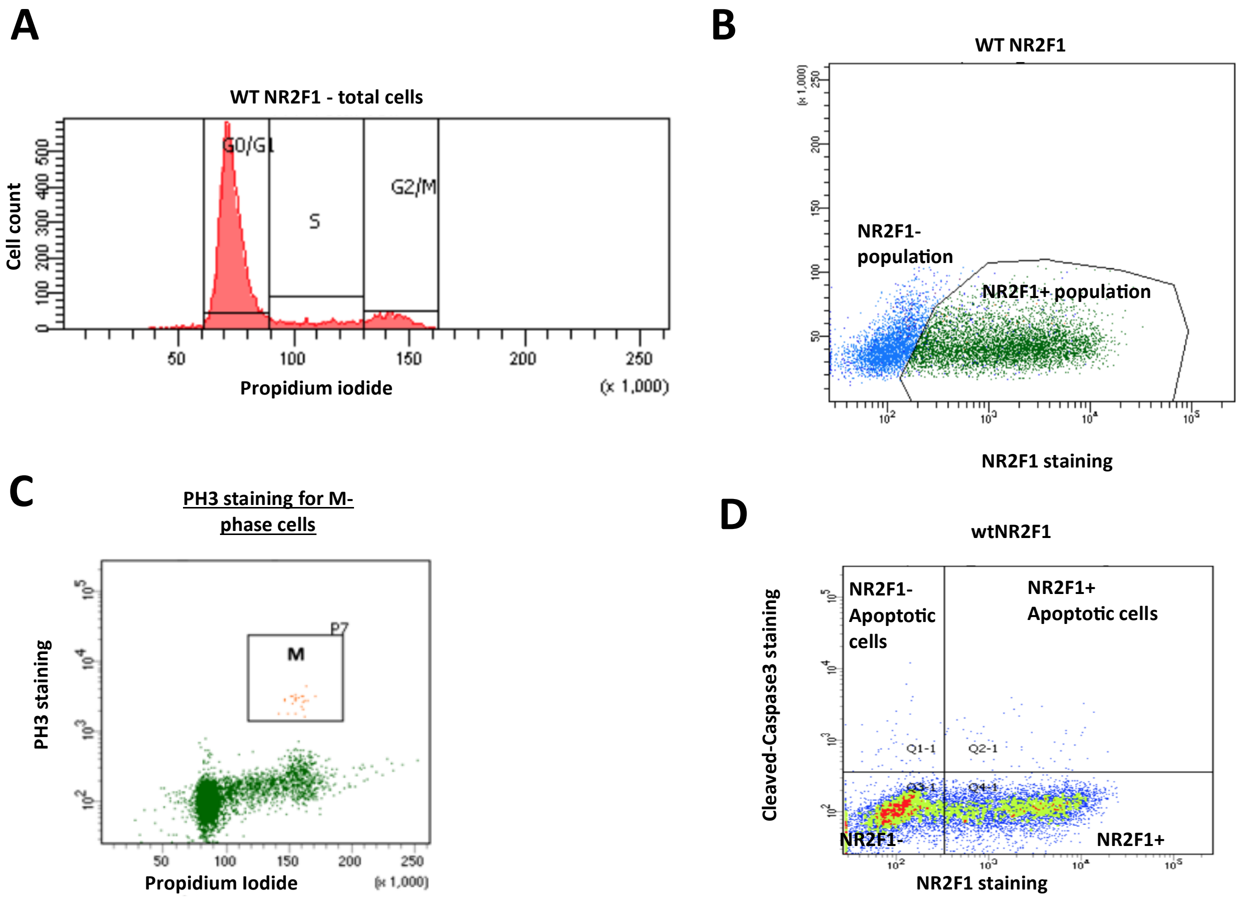


***Figure S3. FACS gating for HEK293T cell cycle and apoptosis analysis.*** *(A) Cell cycle phases (G0/G1, S, G2/M) of NR2F1-WT expressing HEK293T cells were analyzed using propidium iodide staining. This figure shows the number of cells in different cell cycle phases. (B) The NR2F1-positive population (green) is distinguished from the NR2F1-negative population (blue) by NR2F1 staining. (C) Cells in the mitotic phase are indicated in the square (M) by PH3 staining along with propidium iodide staining. (D) Cleaved-caspase 3 was used to indicate apoptotic cells. NR2F1-positive apoptotic cells are cleaved-caspase3 and NR2F1 double-positive and are indicated in the upper right quadrant.*

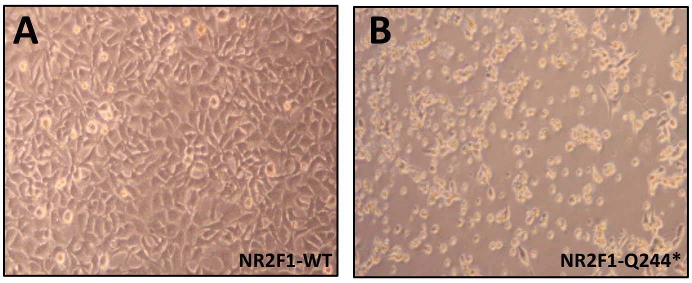


***Figure S4. Morphology of HEK293T cells expressing the truncated NR2F1-Q244* protein**. After 48 hours of transfection of NR2F1-WT (**A**) and NR2F1-Q244* (**B**), cells were observed under a bright-field microscope. The lower number of cells expressing NR2F1-Q244* was observed with a significant portion of cells exhibiting the morphology corresponding to apoptotic cells.*

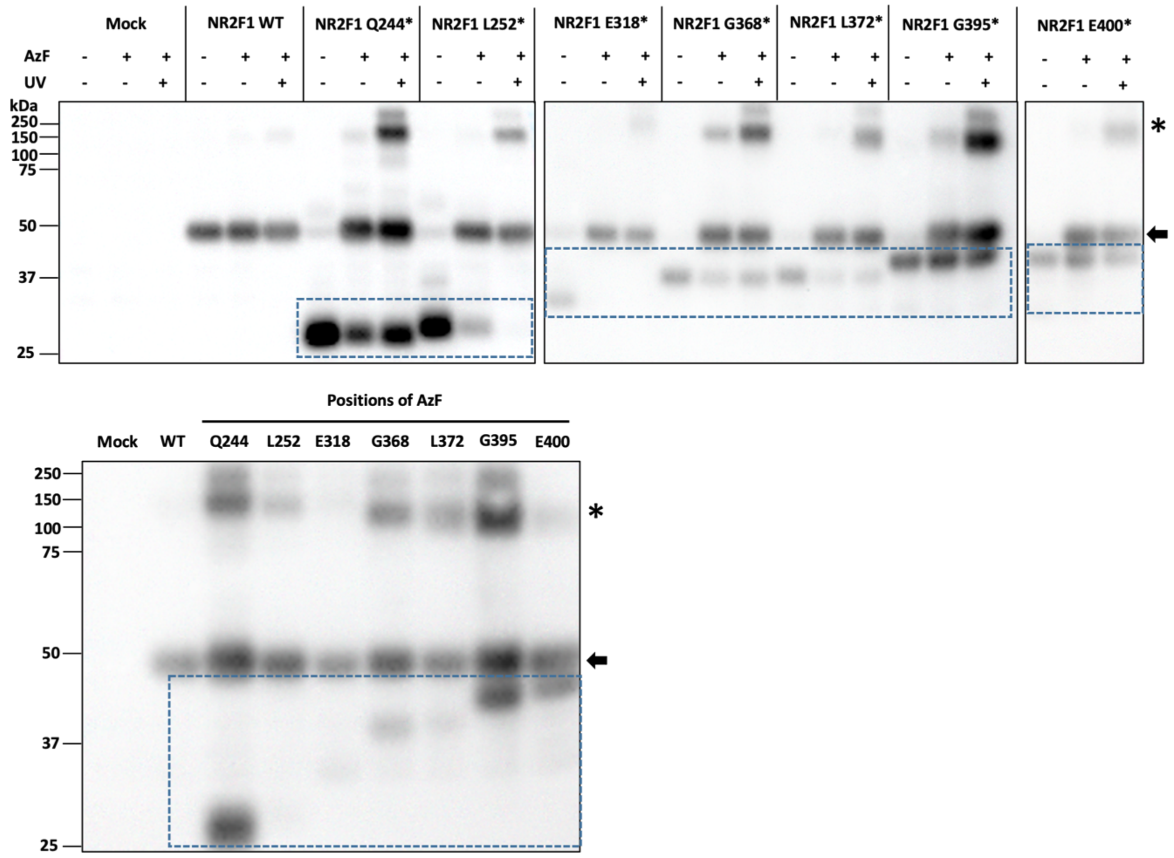


***Figure S5. Site-specific photo-crosslinker incorporation in the LBD of NR2F1 and covalent protein complex capturing in living HEK293T cells.*** *AzF incorporation was performed as described in* ***Figure 6*** *and Methods. Top: Immunoblots show the AzF incorporation after 48 hours of transfection. Truncated NR2F1 (boxed) was produced in the absence of AzF. Once AzF was present in cultural media, the full-length NR2F1 (arrow) was produced by amber codon suppression. The large protein complexes (*) can be observed where NR2F1-AzF protein was produced and photo-crosslinked by UV irradiation (365 nm). Bottom: Immunoblot showing different molecular weights of truncated forms which varied depending on the positions of the amber codon placed in the LBD. The difference in the molecular weights of the photo-crosslinked complexes of NR2F1 proteins that carried AzF at different positions suggests the difference in the protein partner identity and/or in the conformation of the crosslinked complex.*

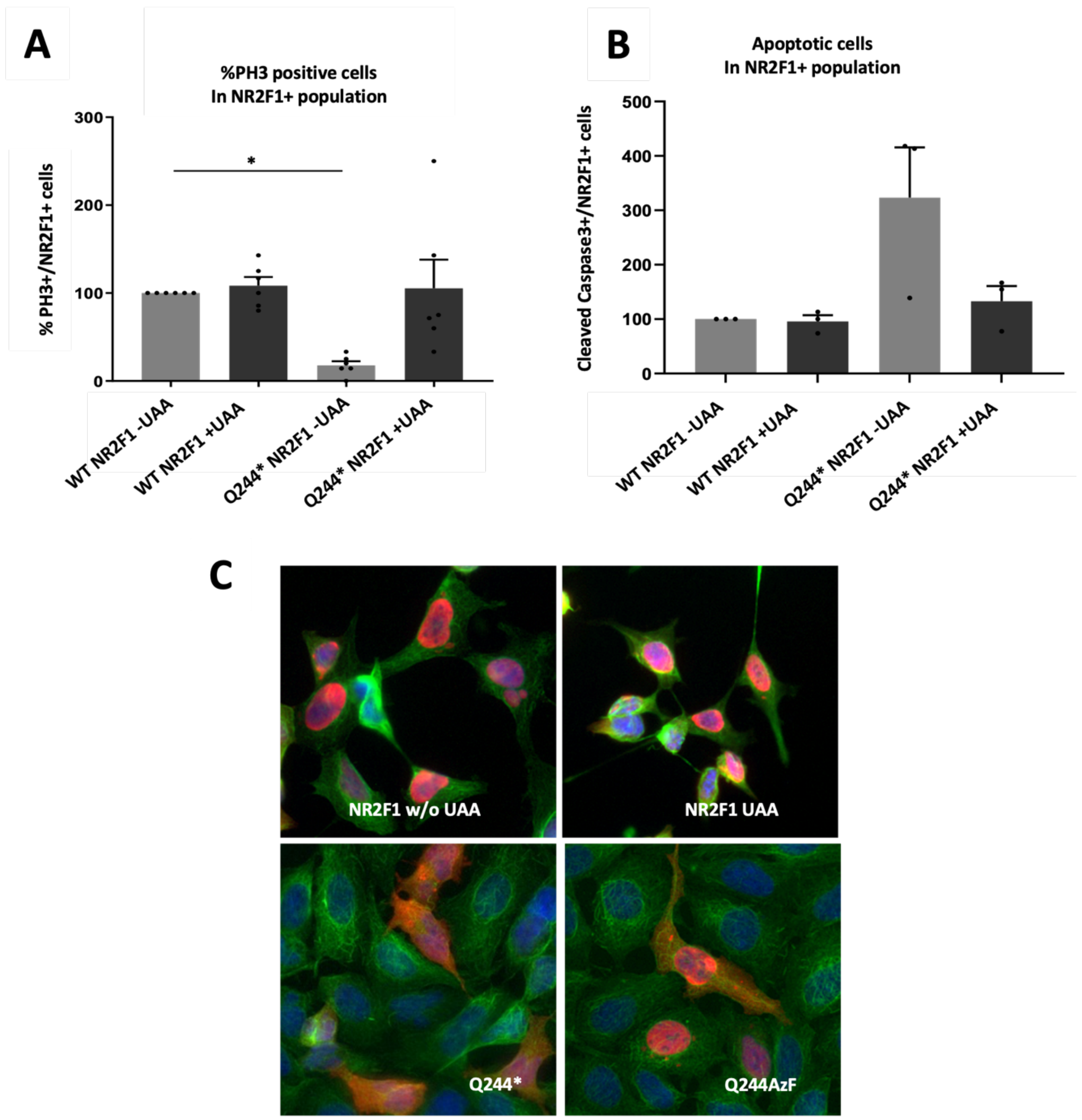


***Figure S6. Full-length NR2F1-Q244AzF functions equivalently to the NR2F1-WT in HEK293T cells.*** *and The quantification of mitotic (**A**) and apoptotic (**B**) cells was carried out by FACS analysis. The comparison between cells expressing the truncated NR2F1-Q244* and the full-length NR2F1-Q244AzF proteins indicated that the incorporation of AzF to yield the full-length NR2F1-Q244AzF restored to the cells the ability to proliferate and rescued the cells from apoptosis to the levels of cells expressing NR2F1-WT. (**C**) Immunostaining of NR2F1 (red) shows that the truncated NR2F1-Q244* localizes in the cytoplasm (co-localization with tubulin, green). The AzF incorporation to yield the full-length NR2F1-Q244AzF protein restores the nuclear localization in the same manner as NR2F1-WT (co-localization with DAPI staining of the nucleus, blue).*

| References | LOVD Database ID; Patient ID | Variant (DNA) | Variant type | Variant (protein) | MRI (general; optic nerve and cortical morphology) | DD | ID | Visual system defect(s) and visual deficit | EOE/S | ASD behavioral abnormalities | Hypotonia | Others |
|---|---|---|---|---|---|---|---|---|---|---|---|---|
| **SA13** | BE18, #6; RE20, #53 | c.1211G>A | *De novo* MM in LBD | **p.Arg404His** | ND | ND | ND | ND | ND | ASD | ND | ND |
| **BO14** | **LOVD: NR2F1_000003**; BO14, #3; BE18, #9; RE20, #48 | c.755T>C | *De novo* MM in LBD | **p.Leu252Pro** | ND | Yes | Yes (IQ 55-65) | P/SOD; CVI; GVI | ND | ND | Yes | ND |
| **CH16** | **LOVD: NR2F1_000078**; CH16, #7; BE18, #22; RE20, #51 | c.1103G>A | *De novo* MM in LBD | **p.Gly368Asp** | Normal or ND | Yes | Yes (IQ ND); speech delay | No or ND | First generalized seizure at 18yo | ASD; RB; aggressive behavior | No | Astigmatism; FD (cupped ears, a small mouth, sloping forehead) |
| **KA17** | **LOVD: NR2F1_000079**; BE18, #37; RE20, #52 | c.1115T>C | *De novo* MM in LBD | **p.Leu372Pro** | ND | Yes, DMD | Yes (IQ ND); speech delay | OA; GVI | ND | RB; ADHD | Yes | Heart murmur. Sleep difficulties. Oromotor disfunction and feeding problems. Motor delay or fine coordination problems / speech delay. |
| **FO19** | **LOVD: NR2F1_000088** | c.603_606del | Frameshift truncation | **p.Arg202Thrfs*154** | ND | ND | ND | ND | ND | ND | ND | ND |
| **FO19** | **LOVD: NR2F1_000019** | c.954G>C | MM in LBD | **p.Glu318Asp** | ND | ND | ND | ND | ND | ND | ND | ND |
| **FO19** | **LOVD: NR2F1_000020** | c.968A>C | MM in LBD | **p.Lys323Thr** | ND | ND | ND | ND | ND | ND | ND | ND |
| **FO19** | **LOVD: NR2F1_000014** | c.1016C>T | MM in LBD | **p.Ala339Val** | ND | ND | ND | ND | ND | ND | ND | ND |
| **FO19** | **LOVD: NR2F1_000025** | c.1025A>G | MM in LBD | **p.Glu342Gly** | ND | ND | ND | ND | ND | ND | ND | ND |
| **FO19** | **LOVD: NR2F1_000021** | c.1117C>T | Truncation | **p.Arg373*** | ND | ND | ND | ND | ND | ND | ND | ND |
| **FO19** | **LOVD: NR2F1_000031** | c.1147_1149del | Deletion in LBD | **p.Ser383del** | ND | ND | ND | ND | ND | ND | ND | ND |
| **FO19** | **LOVD: NR2F1_000031** | c.1147_1149del | Deletion in LBD | **p.Ser383del** | ND | ND | ND | ND | ND | ND | ND | ND |
| **FO19** | **LOVD: NR2F1_000027** | c.1158G>T | MM in LBD | **p.Glu386Asp** | ND | ND | ND | ND | ND | ND | ND | ND |
| **FO19** | **LOVD: NR2F1_000032** | c.1183G>T | MM in LBD | **p.Gly395Cys** | ND | ND | ND | ND | ND | ND | ND | ND |
| **FO19** | **LOVD: NR2F1_000089** | c.1184G>A | MM in LBD | **p.Gly395Asp** | ND | ND | ND | ND | ND | ND | ND | ND |
| **BE20** | **LOVD: NR2F1_000060**; BE20, #2 | c.729_730delinsCT | *De novo* truncation | **p.Gln244*** | CC thinning; ventricular asymmetry and enlargement; abnormal gyration; polymicrogyria | Yes | Yes | No or ND | ND | Behavioral disorders | Yes | Bilateral inguinal hernias. Motor delay or fine coordination problems. |
| **BE20** | **LOVD: NR2F1_000061**; BE20, #6 | c.967_968delAA | *De novo* frameshift truncation | **p.Lys323Serfs*73** | Short CC; ON and chiasm thinning; hypoplastic olfactory lobes; abnormal gyration | Yes, DMD | Yes (speech difficulties) | OA; LVA | ND | ASD traits | No | Pectus excavatum, pes planus, scoliosis, high palate, gastroesophageal reflux, severe apnea, strabismus, hypermetropia, tubular vision. Motor delay. |
| **ZO20** | **LOVD: NR2F1_000085** | c.602C>A | *De novo* truncation | **p.Ser201*** | ND | Mild/moderate | Mild/moderate | Bilateral P/SOD; LVA | ND | ND | ND | Color vision (Ishihara plates) was 1/8 OD and 3/8 OS |
| **WA20** | **LOVD: NR2F1_000051** | c.1080del | Frameshift truncation | **p.Asn362fs*33** | CC; ON and OC hypoplasia; mild MCP | Apparent at 8mo | Speech delay | Severe GVI | Myoclonic astatic seizures at 2½yo | ASD | ND | Phimosis, adenotomy and tympanic drainage. At the age of 5 years, a cervico-thoracic syringomyelia was diagnosed. Hearing defects. |
| **RE20** | **LOVD: NR2F1_000077**; RE20, #49 | c.931G>C | MM in LBD | **p.Ala311Pro** | Normal or ND | Yes | Mild (FSIQ 80 below average); speech delay | P/SOD; mild GVI | Generalized Myoclonic and absence seizures | ASD | Yes | Sleep difficulties. |
| **RE20** | **LOVD: NR2F1_000019**; RE20, #50 | c.954G>C | *De novo* MM in LBD | **p.Glu318Asp** | Abnormal | No but mild DMD | No (IQ 94; performance IQ 54) | OA; CVI; GVI | Atonic; Rolandic epilepsy | ASD | No | Mild/resolved sleep difficulties. High pain tolerance. Mild motor delay or fine coordination problems. |
| **RE20** | **LOVD: NR2F1_000021**; RE20, #46 | c.1117C>T | *De novo* truncation | **p.Arg373*** | CC; ON and OC thinning | Yes, DMD | Mild (DQ ca. 60-70); speech delay | P/SOD; ONH; CVI; GVI | No | ASD | Yes | Touch sensitivity. High pain tolerance. Oromotor disfunction and feeding problems. Motor delay. |
| **RE20** | **LOVD: NR2F1_000036**; RE20, #54 | c.1217T>C | *De novo* MM in LBD | **p.Met406Thr** | Small ON; Abnormal MRI | Yes, DMD | Yes (IQ ND); speech delay | CVI; GVI | No | ASD | Yes | Sleep difficulties. High pain tolerance. Oromotor disfunction and feeding problems. Motor delay. |
| **JS20** | **LOVD: NR2F1_000036** | c.1217T>C | *De novo* MM in LBD | **p.Met406Thr** | DM | Yes | Severe (IQ ND); speech delay; non-verbal | OA; suspected ON dysplasia; GVI | Seizures from 4mo | Short attention span | ND | Recurrent infections; narrow hands with long fingers and dorsal dimpling, narrow foot with long toe and ples planovalgus; Upper and lower limbs contractures; Abnormal pain perception. Motor delay: Gait imbalance, broad-based gait, cerebral palsy. |
| **JU21** | JU21, #11 | c.599 C>G | *De novo* MM in LBD | **p.Thr200Arg** | Lateral and third ventricles enlargement; MCP | Global | Yes (IQ ND); learning disability | Central, steady, maintained | No | ND | Yes | Congenital heart disease: atrial septal defect and patent ductus arteriosus (surgical repair). Chronic lung disease with pulmonary hypertension (bilateral upper pulmonary vein atresia). Congenital bilateral foot deformities: right vertical talus (surgical repair) and mild left hindfoot valgus deformity. Left cryptorchidism (surgical repair). Severe preterm birth (gestational age 27 weeks and 3 days). Non-identical twin has evidence of developmental delay with complications of prematurity. Oromotor disfunction and feeding problems. Motor delay: Balance and coordination deficit. |
| **JU21** | JU21, #12 | c.698G>A | *De novo* truncation | **p.Trp233*** | CC; ON and OC thinning; WM delayed maturation; brain abnormalities | Yes | Yes (IQ ND); speech difficulties | OA; microphthalmia; small ON head; CVI | No | Limited concentration and short attention span | ND | Congenital heart disease: bicuspid aortic valve. Left lacrimal duct stenosis. Recurrent ear infections with left conductive hearing impairment. Microphthalmia. Asymmetric head and alopecia on the crown. Increased tone left ankle. Sister has craniofacial abnormalities (genetically not tested). Motor delay: Toe to heel walking. |
| **JU21** | **LOVD: NR2F1_000082**; JU21, #13 | c.1024G>A | *De novo* MM in LBD | **p.Glu342Lys** | Normal CC e ON; OC atrophy and defective rotation; Normal gyration | No | No | OA; ONH; LVA | No | ND | ND | ND. |
| **JU21** | JU21, #14 | c.1036_1047del | (Likely) *De novo* deletion in LBD | **p.Glu346_Gln349del** | ND | Yes; walking delay | Yes (IQ ND); speech delay; dyslexia; learning disability | ONH; CVI; LVA | No | ND | ND | Frequent ear infections. Cleft palate (surgical repair). Suspected Pierre-Robin sequenced (17q24.3-q25.1 excluded by interrogating WGS data). Asthma. Joint hypermobility syndrome. Periventricular leukodystrophy from premature birth. Motor delay: Balance and coordination deficit. |
| **JU21** | **LOVD: NR2F1_000079**; JU21, #15 | c.1115T>C | Familial MM in LBD | **p.Leu372Pro** | ND | Yes; walking delay | Yes (IQ ND); speech delay | Small ON head; CVI; LVA | No | ND | ND | Motor delay: clumsy. |
| **JU21** | **LOVD: NR2F1_000079**; JU21, #16 | c.1115T>C | Familial MM in LBD | **p.Leu372Pro** | Normal or ND | Yes; walking delay | Yes (IQ ND); speech delay; | OA; ONH; CVI; LVA | One episode of FS | ND | ND | Motor delay: clumsy. |
| **JU21** | **LOVD: NR2F1_000079**; JU21, #17 | c.1115T>C | Familial MM in LBD | **p.Leu372Pro** | Normal or ND | Yes; walking delay | Yes (IQ ND); speech delay; | OA; CVI; LVA | No | ND | ND | ND |
| **JU21** | JU21, #18 | c.1118_1123del | Familial deletion in LBD | **p.Arg373_Leu374del** | ND | No | ND | OA; ONH; LVA | No | ND | ND | ND |
| **JU21** | JU21, #19 | c.1118_1123del | Familial deletion in LBD | **p.Arg373_Leu374del** | ON atrophy | No | ND | OA; ONH; LVA | No | ND | ND | ND |
| **JU21** | JU21, #20 | c.1183G>A | *De novo* MM in LBD | **p.Gly395Ser** | ON atrophy; WM loss | Yes | Yes (IQ ND); learning disability | CVI; LVA | No | ND | Generalized | Motor delay: Balance and coordination deficit. |
| **JU21** | **LOVD: NR2F1_000083**; JU21, #21 | c.1198G>T | *De novo* truncation | **p.Glu400*** | CC thinning; ON and OC atrophy; abnormal gyration | Yes | Yes (IQ ND); learning disability | OA; ONH; LVA | No | ASD; behavioral disorders | ND | Motor delay: fine motor disorder, clumsy, motor dyspraxia. |
| **BI21** | **LOVD: NR2F1_000090** | c.854C>A | Truncation | **p.Ser285*** | ND | ND | ND | ND | ND | ND | ND | ND |
| **BI21**; **STUDER lab** Unpublished | **LOVD: NR2F1_000081** | c.883T>C | MM in LBD | **p.Phe295Leu** | ND | No | No | OA; P/SOD; GVI | ND | ND | ND | Reduced number of RGCs, bilateral and symmetric (but no alterations in RNFL); Reduced visual acuity from 7p2 (right) and 8p2 (left). Papillary pallor in right eye. |
| **BI21** | **LOVD: NR2F1_000091** | c.965T>A | MM in LBD | **p.Leu322His** | ND | ND | ND | ND | ND | ND | ND | ND |
| **BI21** | **LOVD: NR2F1_000087** | c.1168_1170del | Deletion in LBD | **p.Phe390del** | ND | ND | ND | ND | ND | ND | ND | ND |
| **STUDER lab** Unpublished | | c.1184G>C | *De novo* MM in LBD | **p.Gly395Ala** | Normal CC; Abnormal gyration | ND | Yes | GVI | Myoclonia | ASD | Yes | Restricted visual field. |
| **STUDER lab** Unpublished; **FO19** | **LOVD: NR2F1_000026** | c.1096C>T | *De novo* MM in LBD | **p.Arg366Cys** | Mild abnormalities | Yes | Yes (speech delay) | ND | No | ND | ND | Sleep difficulties. Motor delay: global. Intermittent strabismus. MRI: mega cisterna magna abnormalities, without abnormalities of the cerebellum. |

**Abbreviations:**

| | |
|---|---|
| **ADHD** | Attention deficit hyperactivity disorder |
| **ASD** | Autism spectrum disorder |
| **CC** | Corpus Callosum |
| **CVI** | Cortical/Cerebral Visual Impairment |
| **DD** | Developmental delay |
| **DM** | Delayed myelination |
| **DMD** | Delayed motor development/ poor coordination |
| **DQ** | developmental quotient |
| **EOE/S** | Early onset epilepsy/Seizures |
| **FD** | Facial dysmorphism |
| **FS** | Febrile seizures |
| **GVI** | General Visual Impairment |
| **ID** | Intellectual disability |
| **IQ** | developmental quotient |
| **LBD** | Ligand binding domain |
| **LVA** | Low visual acuity |
| **MCP** | macrocephaly |
| **MM** | Missense mutation |
| **MRI** | Magnetic resonance imaging |
| **OA** | Optic atrophy |
| **OC** | optic chiasm |
| **ON** | Optic nerve |
| **ONH** | ON hypoplasia |
| **P/SOD** | Pale/small optic disc |
| **RB** | repetitive behavior |
| **RGCs** | Retinal ganglion cells |
| **RNFL** | Retinal nerve fibre layer |
| **WGS** | Whole genome sequencing |
| **WM** | white matter |

**REFERENCES:**

| | |
|---|---|
| **SA13** | Sanders et al., 2013 |
| **BO14** | Bosch et al., 2014 |
| **CH16** | Chen et al., 2016 |
| **KA17** | Kaiwar et al., 2017 |
| **FO19** | Fokkema et al., 2019 |
| **BE18** | Bertacchi et al., 2018 |
| **BE20** | Bertacchi et al., 2020 |
| **ZO20** | Zou et al., 2020 |
| **WA20** | Walsh et al., 2020 |
| **RE20** | Rech et al., 2020 |
| **JS20** | Jezela-Stanek et al., 2020 |
| **JU21** | Jurkute, Bertacchi et al., 2021 |
| STUDER lab Unpublished | New BBSOAS clinical report (or new clinical description of a |

***Table S1. Summary list of NR2F1 variants in the LBD and clinical description of BBSOAS reported patients.*** *BBSOAS patients, identified by their protein variant and—when available—by their LOVD identifier, are listed following the chronological order of reports and publications describing their cases. Main clinical signs include altered brain morphology as observed by MRI, developmental delay (DD), intellectual disability (ID), visual system deficits, early-onset epilepsy and seizures (EOE/S), autism spectrum disorder (ASD), and behavioral abnormalities and hypotonia. An extended version of these data can be found in Bertacchi et al., 2022. Abbreviations and reference lists are listed below the table.*

| Residue | $\Delta\Delta G_f^{app}$ (kcal/mol) | | | | | $\Delta\Delta G_b^{app}$ (kcal/mol) | | | |
|---|---|---|---|---|---|---|---|---|---|
| | Isolated | NR2F1 (het) | NR2F1 (homo) | NR2F2 | RXRα | NR2F1 (het) | NR2F1 (homo) | NR2F2 | RXRα |
| T200R | -11.1 | -1.4 | -1.5 | -0.8 | -0.01 | -0.4 | -0.7 | -0.4 | 0.3 |
| L252P | 67.5 | 64.3 | 128.5 | 63.8 | 63 | -0.5 | -1.1 | -0.6 | -0.9 |
| F295L | 5.1 | 1 | 1.9 | 1 | 1 | -0.03 | -0.05 | -0.01 | -0.02 |
| A311P | 53.4 | 51.4 | 102.8 | 50.2 | 49.3 | 0.3 | 0.6 | 0.2 | 0.2 |
| E318D | 22.1 | 24.4 | 48.8 | 24.4 | 24.4 | 0.3 | 0.6 | 0.3 | 0.4 |
| E342K | 6.1 | 7.7 | 16.6 | 7 | 18.9 | -0.07 | 1.1 | 0.5 | 16 |
| R366C | 42.4 | 43.1 | 86.4 | 42.9 | 42.9 | 1.6 | 3.4 | 1.6 | 1.5 |
| G368D | -4 | 183.7 | 369.1 | 118.2 | 143.1 | 74.3 | 150.4 | 55.2 | 82.6 |
| L372P | 55.1 | 75.7 | 151.4 | 69.2 | 64 | 6.4 | 13 | 8.7 | 9.3 |
| G395A | 22.6 | 16.7 | 33.3 | 16.7 | 16.7 | 0 | -0.01 | 0 | 0 |
| G395S | 23.8 | 17 | 33.9 | 17 | 17 | -0.005 | -0.01 | -0.01 | -0.01 |
| R404H | 1.4 | 12.7 | 22.4 | 12.8 | 12.4 | -0.1 | -0.2 | 0.07 | -0.4 |
| M406T | 22.2 | 24 | 48.1 | 24.1 | 24.1 | -0.03 | -0.06 | -0.01 | -0.03 |

**Table S2. Predicted effects of BBSOAS-associated variants of NR2F1 LBD on the stability (ΔΔGfapp) of the isolated auto-repressed NR2F1 LBD and on both the stability and affinity (ΔΔGbapp) of the auto-repressed NR2F1 LBD in complex with LBD from NR2F1, NR2F2 and RXRα.** For NR2F1 homodimers, the effects of the mutations were evaluated in both heterozygosis (het) and homozygosis (hom) genetic conditions. ΔΔG values are expressed in kcal/mol and are highlighted in a color scale red-yellow-green from the most to the least detrimental to the respective property. Negative values indicate a stabilizing effect of the mutation.

| Residue | $\Delta\Delta G_f^{app}$ (kcal/mol) | | | | | $\Delta\Delta G_b^{app}$ (kcal/mol) | | | |
|---|---|---|---|---|---|---|---|---|---|
| | Isolated | NR2F1 (het) | NR2F1 (homo) | NR2F2 | RXRα | NR2F1 (het) | NR2F1 (homo) | NR2F2 | RXRα |
| T200R | -4.29 | -3.49 | -15.64 | -3.78 | -3.26 | -0.07 | -0.13 | -0.14 | 0.22 |
| L252P | 55.88 | 55.76 | 111.52 | 56.18 | 55.36 | -0.34 | -0.67 | -0.25 | -0.39 |
| F295L | -4.60 | -0.38 | -0.81 | -0.49 | 0.27 | -0.12 | -0.25 | -0.14 | -0.01 |
| A311P | 55.22 | 52.15 | 104.29 | 51.90 | 52.53 | 0.26 | 0.52 | 0.38 | 0.35 |
| E318D | 23.23 | 23.05 | 46.01 | 22.77 | 23.69 | 0.22 | 0.42 | 0.20 | 0.34 |
| E342K | 3.20 | 18.33 | 37.28 | 7.17 | 12.53 | 12.88 | 26.68 | 1.08 | 5.87 |
| R366C | 54.27 | 53.68 | 107.47 | 54.37 | 54.09 | 1.81 | 3.76 | 2.28 | 1.80 |
| G368D | -3.96 | 228.27 | 422.79 | 171.49 | 295.23 | 131.47 | 229.66 | 64.98 | 140.41 |
| L372P | 56.65 | 74.62 | 149.33 | 68.87 | 70.27 | 10.67 | 21.42 | 8.58 | 10.37 |
| G395A | 1.09 | 2.62 | 2.95 | 1.49 | 1.50 | 0.02 | 0.00 | 0.00 | 0.01 |
| G395S | 0.96 | 1.47 | 6.72 | 2.54 | 2.65 | 0.01 | 0.00 | 0.00 | 0.02 |
| R404H | 9.88 | 7.99 | 15.17 | 7.81 | 7.64 | -0.06 | -0.33 | -0.12 | -0.28 |
| M406T | 24.07 | 23.75 | 47.55 | 23.70 | 23.79 | 0.00 | 0.00 | -0.01 | 0.01 |

***Table S3. Predicted effects of BBSOAS-associated variants of NR2F1 LBD on the stability ($\Delta\Delta G_f^{app}$) of the isolated active NR2F1 LBD and on both the stability and affinity ($\Delta\Delta G_b^{app}$) of the active NR2F1 LBD in complex with LBD from NR2F1, NR2F2 and RXRα.*** *For NR2F1 homodimers, the effects of the mutations were evaluated in both heterozygosis (het) and homozygosis (hom) genetic conditions. ΔΔG values are expressed in kcal/mol and are highlighted in a color scale red-yellow-green from the most to the least detrimental to the respective property. Negative values indicate a stabilizing effect of the mutation.*

| State | Native-like poses[a] | Highest ranked pose[b] | ZDOCK-score[c] |
|---|---|---|---|
| **Auto-repressed** | 23 | 77, 51, 47, 42 | 49.31 ± 3.97 |
| **Active** | 23 | 3, 4, 2, 1 | 58.06 ± 6.63 |

***Table S4. Results from rigid-body docking performed with ZDOCK 2.3.*** *[a]Number of docked complexes with a Cα-RMSD < 1 Å with respect to the native structure obtained by docking simulations performed with PIPER. [b]Rank of the native-like docked pose with the highest score among the 4000 poses of each of the four docking replicas. [c]ZDOCK-score of the native-like solutions reported as average ± standard deviation.*

### Amber codon

| | | |
|---|---|---|
| **Q244TAG** | hCOUP_Q244TAG_f | CTTCCCGGATCTGTAGATCACCGACCAGGTG |
| | hCOUP_Q244TAG_r | CAGATCCGGGAAGAAGGGGATGTTGCGG |
| **L252TAG** | hCOUP_L252TAG_f | GTGTCCCTGTAGCGCCTCACCTGGAGCGAG |
| | hCOUP_L252TAG_r | CGCTACAGGGACACCTGGTCGGTGATCTGCAG |
| **E318TAG** | hCOUP_E318TAG_f | GACTCAGCCTAGTACAGCTGCCTCAAAGCCATCG |
| | hCOUP_E318TAG_r | CTGTACTAGGCTGAGTCGACGTGTAGCGCCTTG |
| **G368TAG** | hCOUP_G368TAG_f | CCGTTTTTAGAAACTGCTGCTGCGACTGC |
| | hCOUP_G368TAG_r | CAGTTTCTAAAAACGGCTGGGCTGGTTGG |
| **L372TAG** | hCOUP_L372TAG_f | CTGTAGCGACTGCCCTCGCTGCGCAC |
| | hCOUP_L372TAG_r | GCAGTCGCTACAGCAGTTTGCCAAAACGGC |
| **G395TAG** | hCOUP_G395TAG_f | CGTTTGGTATAGAAAACCCCCATCGAAACTCTCATCC |
| | hCOUP_G395TAG_r | GGTTTTCTATACCAAACGGACGAAGAAGAGCTGCTC |
| **E400TAG** | hCOUP_E400TAG_f | CCCCATCTAGACTCTCATCCGCGATATGTTACTGTCTG |
| | hCOUP_E400TAG_r | GAGAGTCTAGATGGGGGTTTTACCTACCAAACGGACG |

### Pathogenic mutations

| | | |
|---|---|---|
| **E318D** | E318D_244TAG_f | GACTCAGCCgatTACAGCTGCCTCAAAGCCATCG |
| | E318D_244TAG_r | CTGTAATCGGCTGAGTCGACGTGTAGCGCCTTG |
| **L252P** | L252P_244TAG_f | CTGCCACGCCTCACCTGGAGCGAGCTG |
| | L252P_244TAG_r | GAGGCGTGGCAGGGACACCTGGTCGGTG |
| **G395A** | G395A_f | GTTTGGTAGCTAAAACCCCCATCGAAACTCTCATCCg |
| | G395A_r | GGTTTTAGCTACCAAACGGACGAAGAAGAGCTGC |
| **G368D** | G368D_f | CCGTTTTGACAAACTGCTGCTGCGACTGCCCTC |
| | G368D_r | GCAGTTTGTCAAAACGGCTGGGCTGGTTGGGG |
| **L372P** | L372P_f | GCTGCCGCGACTGCCCTCGCTGCGC |
| | L372P_r | GTCGCGGCAGCAGTTTGCCAAAACGGCTGGG |
| **G395A** | G395A_400TAG_f | GTTTGGTAGCTAAAACCCCCATCTAGACTCTCATCCg |
| | G395A_400TAG_r | GGTTTTAGCTACCAAACGGACGAAGAAGAGCTGC |

***Table S5. List of primers used in this study.***

| Antigen | Manufacturer and reference | | Species | Working dilution | Experimen t |
|---|---|---|---|---|---|
| NR2F1 | Abcam | ab181137 | Rabbit | 1:5000 (WB) 1:1000 (IF) | WB, IF |
| DDDDK tag (FLAG) | Genetex | GTX115043 | Rabbit | 1:1000 | WB |
| ANTI-FLAG® M2 Affinity Gel | Sigma | A2220 | Mouse | - | IP |
| Myc tag | Cell signaling | 2276 | Mouse | 1:1000 | WB |
| phospho-Histone H3 | Millipore | 06-570 | Rabbit | 1:1000 | IF |
| Acetylated-Tubuline | Sigma | - | Mouse | 1:1000 | IF |
| Histone H3 | R&D | MAB9448 | Rabbit | 1:1000 | WB |
| Cleaved Caspase 3 | Cell signaling | 9661 | Rabbit | 1:1000 | IF |
| CRABP 2 | Proteintech | 10225-1-AP | Rabbit | 1:1000 (WB) 1:1000 (IF) | WB, IP, IF |
| Ms IgG – AF488 | Thermofisher | A11029 | Goat | 1:500 | IF |
| Rb IgG – AF 647 | Thermofisher | A32733 | Goat | 1:500 | IF |
| Rb IgG – AF 488 | Thermofisher | A11034 | Goat | 1:500 | IF |
| Ms IgG – AF 647 | Thermofisher | A21236 | Goat | 1:500 | IF |
| Rb IgG – HRP | Cell signaling | #7074 | Goat | 1:5,000 | WB |
| Ms IgG – HRP | Biorad | 172-1012 | Goat | 1:10,000 | WB |
| Ms IgG2A - HRP | Jackson immunoResearch | 115-035-206 | Goat | 1:10,000 | WB |

***Table S6. List of antibodies used in this study.***

***Video S1.*** *Three-dimensional structure of NR2F1-DNA complex. Protein and DNA structures are represented as a cartoon, with the LBD shown in green, the DBD in cyan, and the dsDNA in purple.*

***Video S2.*** *The three-dimensional structure of the LBD is shown as a cyan cartoon with the AF2 helix shown in red and the CRS in green. The Cα of the residues whose mutations are associated with BBSOAS is represented as yellow spheres and labeled.*

***Video S3.*** *Three-dimensional structure of NR2F1 LBD. Protein structure is shown as a cyan cartoon, the AF2 helix is shown in red and the CRS in green. The Cα of the residues whose mutations are associated with BBSOAS is represented as spheres, labeled, and colored in a red-yellow-green scale according to the ∆∆Gfapp values of the auto-repressed NR2F1 homodimer in heterozygosis reported in table ST1.*
